# Genomic patterns of introgression in interspecific populations created by crossing wheat with its wild relative

**DOI:** 10.1101/855106

**Authors:** Moses Nyine, Elina Adhikari, Marshall Clinesmith, Katherine W. Jordan, Allan K. Fritz, Eduard Akhunov

## Abstract

Introgression from wild relatives is a valuable source of novel allelic diversity for breeding. We investigated the genomic patterns of introgression from *Aegilops tauschii*, the diploid ancestor of the wheat D genome, into winter wheat (*Triticum aestivum*) cultivars. The population of 351 BC_1_F_3_:_5_ lines was selected based on phenology from crosses between six hexaploid wheat lines and 21 wheat-*Ae. tauschii* octoploids. SNP markers developed for this population and a diverse panel of 116 *Ae. tauschii* accessions by complexity-reduced genome sequencing were used to detect introgression based on the identity-by-descent analysis. Overall, introgression frequency positively correlated with recombination rate, with a high incidence of introgression at the ends of chromosomes and low in the pericentromeric regions, and was negatively related to sequence divergence between the parental genomes. Reduced introgression in the pericentromeric low-recombining regions spans nearly 2/3 of each chromosome arm, suggestive of the polygenic nature of introgression barriers that could be associated with multilocus negative epistasis between the alleles of wild and cultivated wheat. On the contrary, negative selection against the wild allele of *Tg*, controlling free-threshing trait and located in the high-recombining chromosomal region, led to reduced introgression only within ∼10 Mbp region around *Tg*. These results are consistent with the effect of selection on linked variation described by the Hill-Robertson effect, and offer insights into the introgression population development for crop imrpovement to ensure retention of introgressed diversity across entire genome.

## Introduction

Wild relatives of wheat show a broad range of adaptation to diverse climatic and biotic stress factors, and for decades served as a source of beneficial allelic diversity in breeding programs for improving drought tolerance (Peleg *et al.* 2005; Reynolds *et al.* 2007), disease resistance (Huang *et al.* 2003; Periyannan *et al.* 2013; Saintenac, Zhang, *et al.* 2013), grain quality (Uauy *et al.* 2006) and yield potential (Cooper *et al.* 2012; Reynolds *et al.* 2017). The availability of high-quality genome assemblies for a number of wild relatives (Ling *et al.* 2013; Avni *et al.* 2017; Luo *et al.* 2017) combined with inexpensive next-generation sequencing technologies allowing for quick genome-wide genotyping of large populations stimulated substantial interest in introgression of the wild relatives’ diversity as a strategy for crop improvement. Allohexaploid bread wheat, *Triticum aestivum* (2n = 6x = 42, genome formula AABBDD), resulted from hybridization between tetraploid *Triticum turgidum* (2n = 4x = 28, genome formula AABB) and *Aegilops tauschii* (2n = 2x = 14, genome formula DD) (Kihara 1944; Luo *et al.* 2007; Wang *et al.* 2013). These two wild relatives serve as the primary sources of allelic diversity for introgression into common wheat (Gill and Raupp 1987; Qi *et al.* 2007; Reynolds *et al.* 2007; Dreisigacker *et al.* 2008). Introgression from these species, which share homologous genomes with wheat, could be easily achieved by either direct crosses between wheat and wild relatives, or crosses between wheat and synthetic wheat lines produced by hybridizing tetraploid wheat and *Ae. tauschii* (Qi *et al.* 2007; Ogbonnaya *et al.* 2013). The wild diploid and tetraploid relatives of wheat that carry homoeologous genomes, such as *T. monococcum* (A^m^ genome), *T. urartu* (A^u^ genome), *T. timopheevii* (2n = 4x = 28, AAGG) or *Ae. speltoides* (2n = 2x = 14, SS) are considered as secondary sources of genetic diversity for introgression. Effective introgression from these species requires recombining the homoeologous chromosomes of common wheat and wild relatives in the absence of *Ph1* gene that controls pairing between homoeologs (Sears 1977).

The retention of wild relative introgression in the populations of cultivated crops could be affected by selection for fitness or traits related to agronomic performance. There was evidence for reduced historic introgression from wild relatives into wheat and maize cultivars around genes controlling major domestication traits (Wang *et al.* 2017; He *et al.* 2019). For example, in wheat, the region of the genome harboring domestication gene *Q* (Simons *et al.* 2006) showed reduced genetic diversity and no evidence of introgression from sympatric populations of wild emmer wheat (He *et al.* 2019), consistent with selection against alleles affecting the domestication phenotype. Interaction between introgressed alleles and the genetic backgrounds of recipient lines could result in hybrid sterility, or influence the general phenology or agronomic performance resulting in the loss or retention of introgressed genome segments (Rieseberg *et al.* 1999; Jiang *et al.* 2000). In sunflower, variation in the frequency and distribution of introgression across genome was attributed to reduced introgression around the structurally re-arranged regions and alleles linked with pollen sterility (Rieseberg *et al.* 1999). A multilocus epistatic interaction was proposed as a possible explanation for restricted introgression in the interspecific populations of polyploid cotton (Jiang *et al.* 2000). Selection on the introgressed variants is also expected to affect linked variation, with the size of the affected region depending on the distribution of recombination rate across genome (Hill and Robertson 1966; Martin *et al.* 2019). Therefore, from the practical perspective, understanding the genomic patterns of wild relative introgression into cultivated crops will be crucial for the effective deployment of novel diversity for crop improvement.

Genotyping approaches based on next-generation sequencing of complexity-reduced genomic libraries substantially accelerated analysis of genetic diversity in large crop genomes (Elshire *et al.* 2011; Saintenac, Jiang, *et al.* 2011, 2013; Poland *et al.* 2012; Jordan *et al.* 2015, 2018). The high proportion of missing data in the low-coverage sequencing datasets can be compensated by genotype imputation using the recently developed reference genome (The International Wheat Genome Sequencing Consortium (IWGSC) 2018). Imputation of ungenotyped SNP markers from a reference panel into a target population takes advantage of regions of identity-by-descent (IBD), thus allowing for the interpolation of SNPs into the target population (Browning and Browning 2013). The power and resolution of association studies have been shown to improve after imputation (Browning and Browning 2012; Jordan *et al.* 2015; Nyine *et al.* 2019).

In this study, we developed a population of winter wheat lines carrying introgression from a diverse set of *Ae. tauschii* accessions selected to represent both the broad genetic and geographic diversity of the species. The boundaries of the introgressed segments in the wheat genome were inferred using the IBD analyses based on the SNP datasets generated by complexity-reduced sequencing of 378 samples including 21 *Ae. tauschii* accessions, 6 hexaploid wheat lines and 351 introgression lines. The ungenotyped D-genome SNPs in the introgression population were imputed from a reference panel of 116 *Ae. tauschii* accessions to improve IBD detection. The distribution of introgression across the genome was investigated to assess its overall effect on genetic diversity, and to evaluate the impact of recombination rate variation and early selection for uniform phenological and developmental characteristics on the introgression frequency in different parts of the wheat genome. The effect of phenotypic selection against a non-adaptive allele contributed by *Ae. tauschii* was investigated around the *Tg* gene on chromosome arm 2DS, which controls tenacious glume trait affecting grain threshability (Sood *et al.* 2009).

## Materials and methods

The study population consisted of 351 BC_1_F_3_:_5_ *Ae. tauschii* introgression lines developed by crossing synthetic *Ae. tauschii*-wheat octoploid lines with hexaploid wheat recurrent parents. The octoploid lines were developed by crossing five hexaploid wheat parents with 21 *Ae. tauschii* accessions (Supporting Information Table S1, File S1). The resulting F_1_ hybrid plants regenerated from rescued embryos were treated with colchicine to produce the synthetic octoploids (Dale *et al.* 2017). The synthetic octoploids were backcrossed once to the respective hexaploid wheat parents or to another wheat line. The BC_1_F_1_ plants were self-pollinated and advanced by single seed descent to the BC_1_F_3_ generation. Seeds from individual BC_1_F_3_ plants were bulked and grown in single rows in the field at the Kansas State University Ashland Research Farm near Manhattan, KS in the 2016-17 growing season. A total of thirty-one families were represented in this experiment. The number of lines per family ranged from 42 to 137, and resulted in a total of 2,861 lines that were planted. The 351 lines used in our study were selected in 2017 from this set of lines. Selection criteria included production of sufficient seed to allow yield testing, general fitness, threshability to allow mechanical harvest and phenology similar to the elite hexaploid parent(s). In addition, 116 diverse *Ae. tauschii* accessions representing *Ae. tauschii* ssp. *tauschii* and *Ae. tauschii* ssp. *strangulata* from different geographical locations were used as the reference panel to impute ungenotyped SNPs in the study population to improve detection of region identical-by-descent (IBD) (Supporting Information Table S2).

### Sequencing complexity-reduced genomic libraries

DNA from the *Ae. tauschii* introgression population and the reference panel samples was extracted using DNeasy 96 Plant DNA extraction kit (Qiagen) following the manufacturer’s protocol. The quality and concentration of the DNA was assessed using PicoGreen® dsDNA assay kit (Life Technologies). Input DNA was normalized to 400 ng (20ul of 20ng/ul) using Qiagility robot (Qiagen). Genotyping by sequencing (GBS) libraries were constructed using the protocol described by Saintenac *et al.* (2013), and then subjected to size selection using the Pippin Prep system (Sage Scientific) to enrich for 270-330 bp fragments. In total, five libraries were produced, representing 80 barcoded accessions each. Each library was sequenced on Illumina NextSeq 500 using a 1 x 75 bp kit for the introgression lines and 1 x 100 bp kit for the reference panel following the Illumina protocol. TASSEL 5.0 GBS v2 pipeline (Glaubitz *et al.* 2014) was used to generate SNPs from the fastq files of the introgression lines and the reference panel. In brief, the raw GBS sequence reads were aligned to the Chinese Spring reference sequence v1.0 (The International Wheat Genome Sequencing Consortium (IWGSC) 2018) using Burrow’s Wheeler Alignment (BWA) software v0.7.17. TASSEL 5.0 GBS v2 default parameters were used in all steps (Glaubitz *et al.* 2014), except the KmerLength for the introgression population was set at 35 to account for shorter read length.

### SNP genotyping and imputation

SNPs for the reference *Ae. tauschii* panel with minor allele frequency (MAF) less than 0.02 and maximum missingness greater than 50% were filtered out using vcf-filter tools v0.1.13. The missing SNPs were imputed using the program Beagle v.5.0 (Browning and Browning 2013) with default parameters (File S2). SNPs from *Ae. tauschii* derived introgression lines were filtered in two steps. First, SNPs from all sub-genomes (A, B and D) with minor allele frequency (MAF) less than 0.05 and maximum missingness greater than 30% were filtered out using vcf-filter tools v0.1.13. The missing SNPs were imputed using the program Beagle v.5.0 with default parameters. In the next step, all A and B genome SNPs, and D genome SNPs with MAF less than 0.01 were excluded from the raw vcf file using vcf-filter tools v0.1.13. The program conform-gt (https://faculty.washington.edu/browning/conform-gt.html) was used to check the concordance of D genome SNP positions between the introgression population and the reference panel based on the Chinese Spring genome v1.0 coordinates (IWGSC, 2018). Missing and ungenotyped SNPs in the D genome of the introgression population were imputed from the reference panel using Beagle v.5.0 (File S3).

### Principal component analysis (PCA)

The population structure of the diverse *Ae. tauschii* accessions and the introgression population was analyzed using the 27,880 D genome SNPs segregating in both populations. The SNP dataset was converted to the hapmap format and imported into TASSEL v.5.0, which was used to calculate the principal components. The first two components were plotted to show the distribution and clustering of the reference panel accessions in relation to the 21 parental *Ae. tauschii* accessions and the entire introgression population. In addition, a total of 13,970 SNPs (File S4), including 4,039, 4,255, 5,222 and 454 from A, B, D genomes and unanchored scaffolds, respectively, were used to evaluate the distribution of *Ae. tauschii*-derived introgression lines on the first two principal components using the wheat parents as grouping factors.

### Genetic diversity

To evaluate the effect of introgression on SNP diversity, the mean number of base differences for each D genome SNP site in all pairwise comparisons (π) among *Ae. tauschii* accessions, introgression lines, and hexaploid wheat lines were calculated using vcftools v0.1.13 and summarized using basic R functions (R Development Core Team 2011). The π values for each chromosome were interpolated by calculating the average of ten SNPs sliding forward by two SNPs using a Perl script and plotted using R package ‘ggplot2’ v3.2.1. The set of D genome SNPs used in this analysis was 5,222 that were retained after filtering out SNPs with minor allele frequency less than 0.05 and maximum missingness greater than 30 % (File S4).

### Recombination hotspots

The distribution of recombination hotspots was analyzed using the imputed D-genome SNPs split into subsets based on families. A combination of custom Perl and R scripts (Nyine *et al.* 2018), were used to convert the SNP alleles to 0, 1, and 2, of which, 0 is homozygous major allele, 1 is heterozygous and 2 is homozygous minor allele. Regions containing monomorphic SNPs were eliminated by the R script. A total of 16 families each having at least 10 progenies plus the respective parents were used in this analysis. A separate custom Perl script was used to count the number of allele phase transitions in each chromosome per individual and recode the flanking SNP positions as break points (Jordan *et al.* 2018). The number of recombination breakpoints (RBP) per 2 Mb sliding window with 1 Mb step size in each chromosome per family was obtained using bedmap option from BEDOPS v2.4.35 (Neph *et al.* 2012). The total RBP per 2 Mbp window across the 16 families was determined and the windows within the 95^th^ percentile were considered as recombination hotspots. The centromere position on each chromosome was based on the Chinese Spring reference genome v1.0 (The International Wheat Genome Sequencing Consortium (IWGSC) 2018; Su *et al.* 2019). Kruskal Wallis test was used to test for significant differences in the distribution of recombination breakpoints in each family.

In order to investigate the effect of sequence divergence (SD) and structural re-arrangements on recombination, we compared hexaploid wheat (Chinese Spring) and the diploid relative, *Ae. tauschii* ssp. *strangulata* (AL8/79) D genomes at the protein level. High confidence D genome gene protein sequences from Chinese Spring v.1.0 and *Ae. Tauschii* v.4.0 (Luo *et al.* 2017) were used. The annotation of the *Ae. tauschii* genome was downloaded from http://aegilops.wheat.ucdavis.edu/ATGSP/annotation/. Local protein BLAST databases were created for each dataset using BLAST+ v2.7.1. Reciprocal blastp was performed between the two species’ genome proteins using default parameters. A Perl script was used to filter out blast hits with percent identity less than 95 and gap opens greater than 0. A file consisting of species chromosome identity, gene name, gene start and end positions was generated from the respective gff3 file. MCScanX software (Wang *et al.* 2012) was used to generate the dot plot and dual synteny plot that were used to compare the structural differences between the genome of *T. aesitvum* and *Ae. tauschii*.

The difference in recombination rate between lines carrying introgression from *Ae. tauschii* ssp. *strangulata* or *Ae. tauschii* ssp. *tauschii* was assessed by comparing the sequence divergence (SD) of the parents with introgression frequency (IF) and the total RBPs estimated from the progenies in six introgression families. A total of 5,222 high stringency filtered D genome SNPs were used to determine the RBP and SD while IBD was used to infer introgression frequency in 5 Mbp chromosome windows. SD in our study corresponds to the average number of polymorphic sites between *Ae. tauschii* and hexaploid wheat parents within 5 Mbp windows. Chromosomes were divided into 2/3 pericentromeric and distal regions. Mann-Whitney U test was used to determine if significant difference in SD, IF and RBP existed between ssp. *strangulata* and ssp. *tauschii-*derived families.

### Identity by Descent detection (IBD)

Introgression of *Ae. tauschii* genome in hexaploid wheat was inferred using IBD. The D genome SNPs from each chromosome were separated and used as input genotype (gt) data for IBD detection. The program Beagle v.4.1 was used to detect IBD segments between introgression lines, hexaploid wheat and *Ae. tauschii* parents using default parameters. The R-package ggplot2 v3.2.1 was used to generate a density plot of IBD segment start per chromosome to show the distribution pattern. All chromosomes were scaled by dividing the IBD values by the individual chromosome length and then multiplied by 100.

The efficiency of introgression was estimated as the percentage of observed proportion of *Ae. tauschii* genome (introgression frequency) in the introgression lines as inferred by IBD to the expected proportion of *Ae. tauschii* in BC_1_F_3:5_. Assuming that recombination events between *Ae. tauschii* and hexaploid wheat D genomes occurred normally in each chromosome, the expected proportion of *Ae. tauschii* genome in the BC_1_F_3:5_ introgression lines was approximated at 25%. The observed proportion of introgression was obtained by dividing the total length of IBD segments from *Ae. tauschii* shared with each line by the genome size of *Ae. tauschii* (4.3 Gb) and multiplied by 100. The result was divided by 25 and multiplied by 100 to get the percentage introgression efficiency. The average, standard deviation, minimum and maximum IBD length shared between introgression lines, introgression lines and hexaploid wheat, introgression lines and *Ae. tauschii* parents were determined, and divided by the chromosome size.

The relationship between IBD and the *Tg* gene on chromosome arm 2DS was explored. The IBD count per 1 kb sliding window was used to compare the frequency of introgression in the *Tg* region. Genes within the *Tg* region (21.8 Mbp to 23.3 Mbp) and their functional annotation were extracted from the Chinese Spring reference gene annotation file. Introgression lines were phenotyped for the tenacious glume trait. The results were used to confirm the presence or absence of wild type alleles depending on whether the introgression segment spanned the *Tg* gene region or not. Genome-wide association analysis of tenacious glume trait with the 27,880 SNP markers was done using GAPIT function in R. A mixed linear model was used and the population structure was controlled using the first three principal components calculated from the markers. A Manhattan plot of negative log_10_ of false discovery rate (FDR) transformed P-values from the D chromosomes was generated in R using ‘qqman’ package.

### Data availability

All supplemental material and relevant data are available at FigShare. Raw sequence reads were deposited at NCBI’s Sequence Read Archive (SRA) under BioProjectID PRJNA637316 and PRJNA637416 and will be accessible upon release through the links below.

https://www.ncbi.nlm.nih.gov/sra/PRJNA637316

https://www.ncbi.nlm.nih.gov/sra/PRJNA637416

## Results

### Genotyping and SNP imputation

A total of ∼315 million high-quality NGS reads with barcodes were generated with an average of ∼2.8 million reads per sample from the diverse *Ae. tauschii* accessions (Supporting Information Table S2). Eighty-eight percent (88%) of the reads were aligned to the Chinese Spring reference sequence v.1.0 (The International Wheat Genome Sequencing Consortium (IWGSC) 2018) with an average of ∼2.4 million reads per sample. The number of SNP sites generated from the TASSEL v. 5.0 GBS v.2 pipeline was 148,430. After filtering out SNPs with MAF less than 0.02, and maximum missingness greater than 50%, the number of retained SNPs was 96,056.

Similarly, ∼1.1 billion high quality reads with barcodes were generated from the introgression population, with an average of ∼3.0 million reads per sample (Supporting Information Table S1). Ninety-six percent (96%) of the reads were aligned to the Chinese Spring reference v1.0 with an average of ∼2.9 million reads per sample. Samples with less than 91,000 reads were excluded from downstream analysis. The number of unfiltered SNPs generated by the TASSEL v.5.0 GBS v.2 pipeline was 281,846. A total of 13,970 SNPs from the A, B, D genomes and unanchored scaffolds were retained after filtering out SNPs with MAF less than 0.05 and maximum missingness greater than 30%. The number of SNPs from the D genome was 37.4 % of the filtered SNP dataset. A second round of filtering was performed on the D genome SNPs to remove sites with MAF less than 0.01 resulting in 142,740 SNPs, out of which, 27,880 also segregated in the diverse set of *Ae. tauschii* accessions (henceforth, reference panel). Using the program Beagle v.5.0 (Browning and Browning 2013), 85,752 SNPs were imputed from the reference panel into the *Ae. tauschii*-derived introgression population.

### Principle component analysis

We used the analysis of principal components to assign the *Ae. tauschii* accessions used as parents for introgression population development to the previously identified lineages (Wang *et al.* 2013). The 137 *Ae. tauschii* accessions (116 reference panel plus 21 parental accessions) formed three distinct clusters when the first two PCs calculated from 27,880 SNPs were plotted (Fig. S1), with 79.2 % of variance explained by the first two PCs. One of the three clusters included accessions known to belong to *Ae. tauschii* ssp. *strangulata* or lineage 2 (L2) (Wang *et al.* 2013). The remaining two clusters included accessions that belong to *Ae. tauschii* ssp. *tauschii* or lineage 1 (L1a and L1b). Cluster L1a was the most heterogeneous one with accessions coming from Afghanistan (AFG), Turkmenistan (TKM), Iran (IRN), Pakistan (PAK) and Tajikistan (TJK) (Table S3). Fifteen of the *Ae. tauschii* parents used to generate the introgression population were assigned to this cluster. More than two thirds of the accessions in cluster L1b were from Turkey (TUR) with only a few accessions from Armenia (ARM), IRN, TJK and PAK. Three parents of the introgression population were present in this cluster. Cluster L2 consisted of *Ae. tauschii* accessions mostly collected from Iran (IRN), although a few accessions from Azerbaijan (AZE), Turkmenistan (TKM), Syria (SYR) and TUR were present. Three parents of the introgression population parents clustered in this group and two of them (TA1642, TA2378) are known to belong to *Ae. tauschii* ssp. *strangulata* or lineage 2 (Wang *et al.* 2013; Singh *et al.* 2019).

When the introgression lines were plotted on the first two PCs together with *Ae. tauschii* accessions and hexaploid wheat parents, clusters L1a and L1b were less distinguishable from each other (Fig. S2). The total variance explained by the first two PCs was 56.7%. The cluster including *Ae. tauschii* accessions from lineage L2 was located in close proximity to cluster including hexaploid wheat parents (HW) consistent with earlier studies demonstrating the origin of the wheat D genome from the *Ae. tauschii* ssp. *strangulata* lineage (Dvorak *et al.* 1998; Wang *et al.* 2013). The introgression lines did not form a clearly distinguishable cluster, and were scattered broadly, indicating a wide range of variation in the proportion of the genome introgressed from the different lineages of *Ae. tauschii* (Fig. S2). Many introgression lines clustered closer to the hexaploid wheat parents indicating that the greater proportion of genome in the BC_1_F_3:5_ lines comes from hexaploid wheat. This trend is likely associated with the loss of the introgressed segments as a result of backcrossing to the hexaploid parents and selection during population development. When the introgression lines were compared with the hexaploid wheat parents using 13,970 SNPs from all three sub-genomes, clustering was consistent with the pedigree (Fig. 1), and co-clustering of introgression lines from different families was observed because of the shared *Ae. tauschii* parents.

**Fig. 1.**
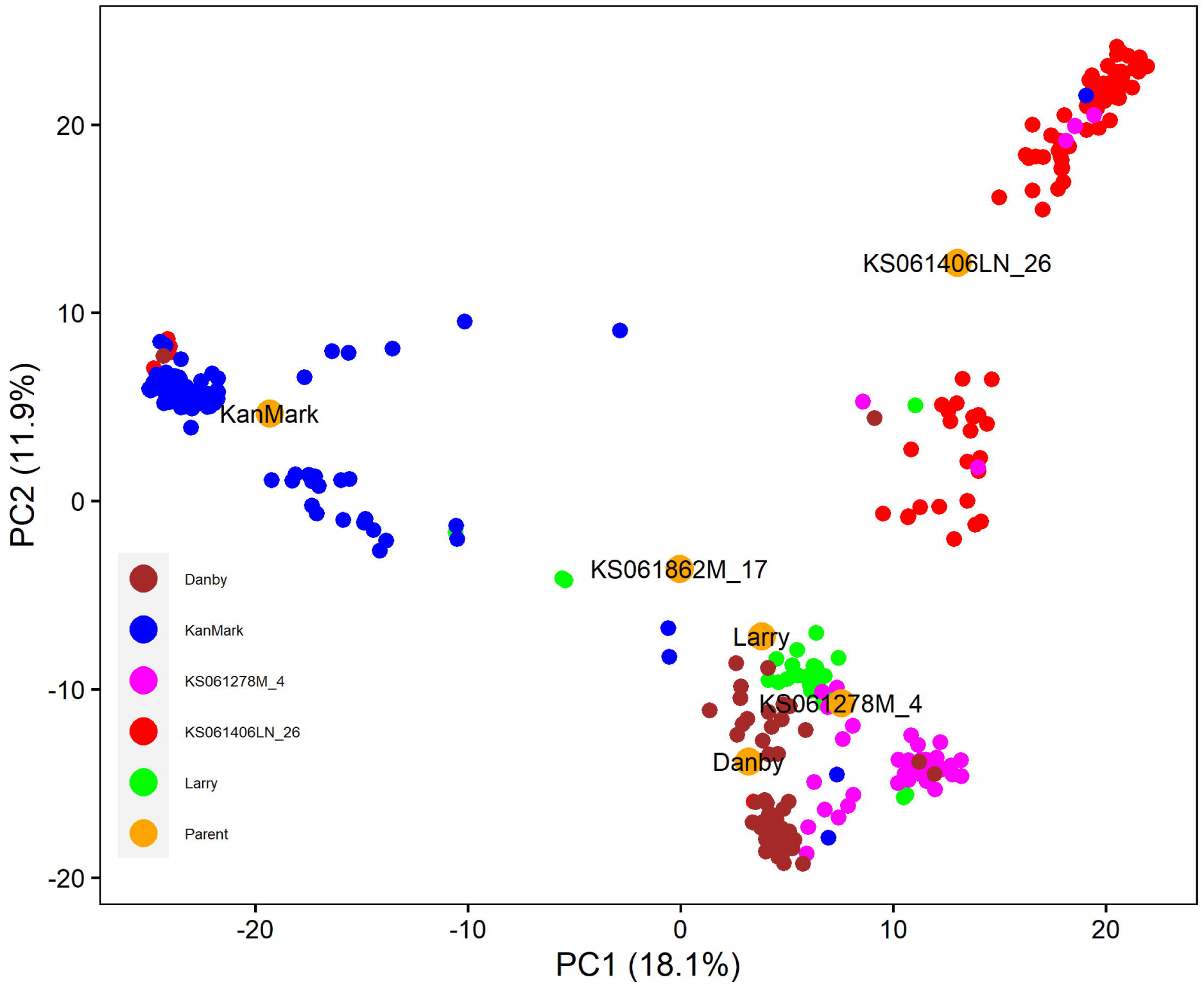
Distribution of the introgression lines and the hexaploid wheat parents on the first two principal components based on SNP markers from A, B, D genomes and unanchored scaffolds.

### Distribution of SNP diversity in the introgression population

To assess the effect of wild relative introgression on genetic diversity in wheat, we estimated SNP diversity (π) in the populations of *Ae. tauschii*, hexaploid wheat parents, and the introgression lines. The average per-site estimate of π for the diverse *Ae. Tauschii* populations was 0.304. Consistent with previous studies (Akhunov *et al.* 2010; Jordan *et al.* 2015; Balfourier *et al.* 2019; Pont *et al.* 2019), a cross-population diversity comparison showed a low average SNP diversity in the wheat D genome across all chromosomes (Table 1). The lowest diversity was found in the hexaploid wheat parents with the chromosome mean ranging from 0.003 to 0.012 as compared to *Ae. tauschii* parents that ranged from 0.118 to 0.131. For most chromosome regions, the levels of SNP diversity in the introgression population were intermediate between the levels of diversity in the parental populations of wheat and *Ae. tauschii*, but tended towards the *Ae. tauschii* with maximum mean π of 0.122 on chromosome 4D (Fig. 2 and Fig. S3). Analysis of variance showed significant differences in π values between *Ae. tauschii*, hexaploid wheat, and introgression lines, and between chromosomes (P < 0.001). The SNP diversity of the introgression lines for most regions of chromosome 4D and 5D were higher than those of *Ae. tauschii* parents (Fig. S3). Taken together, these results indicate that *Ae. tauschii* introgression substantially increased the SNP diversity of the recurrent hexaploid wheat parents.

**Table 1.**
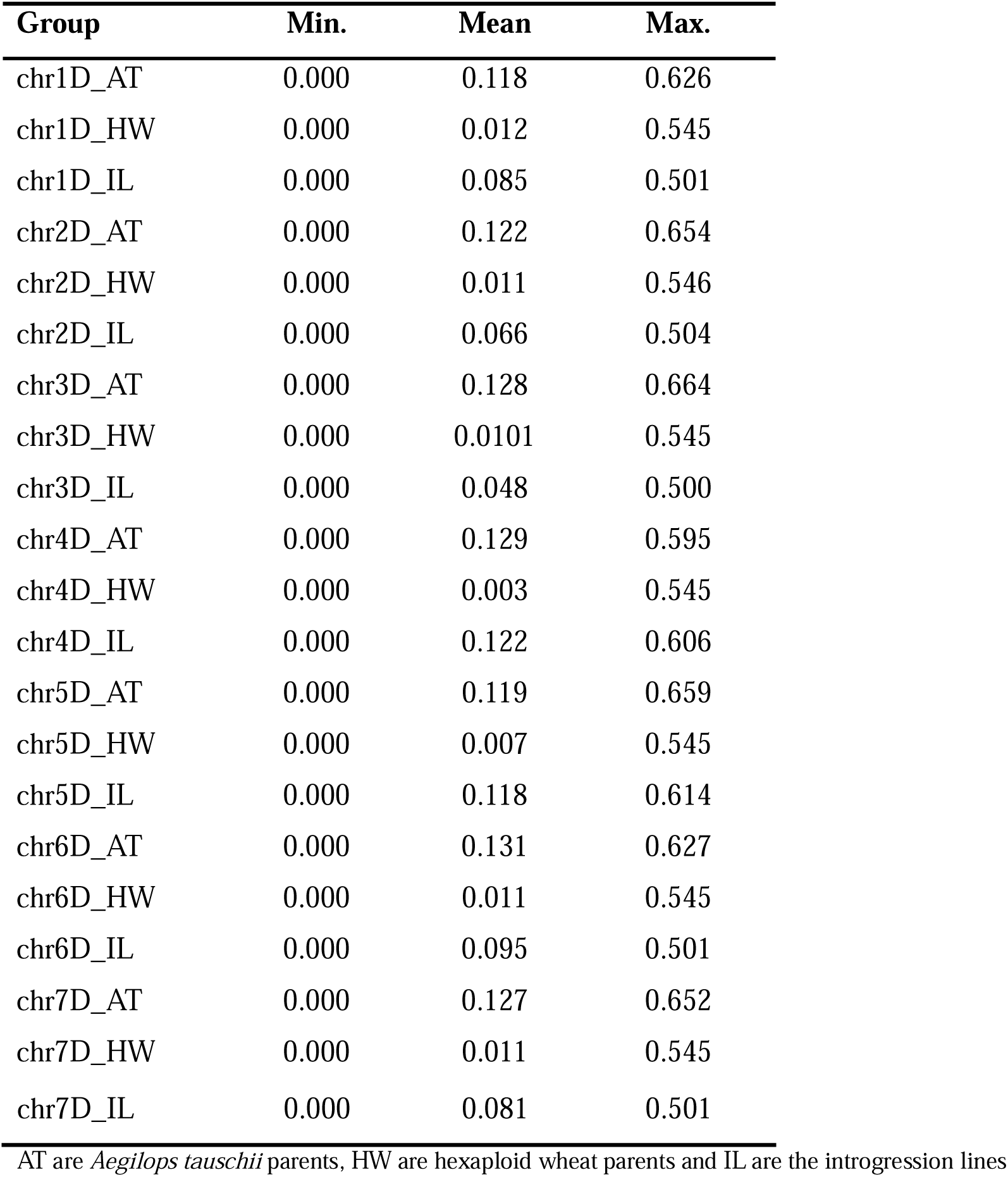
A comparison of nucleotide diversity of *Ae. tauschii* derived introgression lines and their parents.

**Fig. 2.**
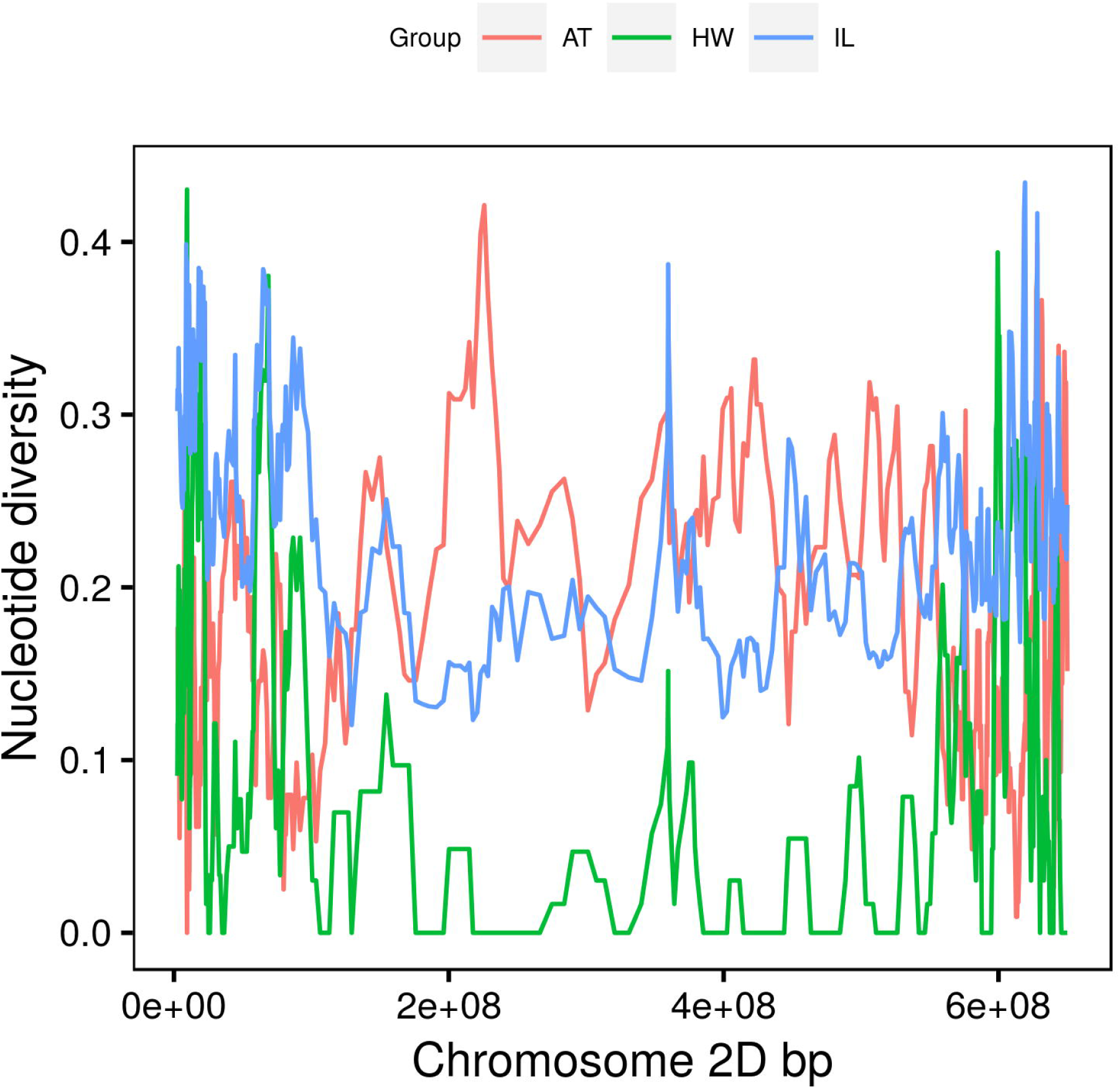
Variation in nucleotide diversity for chromosome 2D based on per estimates of π for *Ae. tauschii* accessions (AT), hexaploid wheat (HW) lines and introgression lines (IL).

### Genomic patterns of introgression and recombination rate

One of the factors affecting the frequency of introgression across the genome are structural re-arrangements (Rieseberg *et al.* 1995, 1999). Comparative analysis of gene order along the chromosomes showed that more than 99% of the genes from *T. aestivum* are perfectly collinear to those of *Ae. tauschii* ssp. *strangulata* suggesting a lack of major structural re-arrangements (Fig. 3). Only small-scale inversions were observed on chromosomes 2D, 4D and 6D in the regions near the centromeres, and four genes were found in non-syntenic positions between the wheat (1D and 5D) and *Ae. tauschii* (1D, 4D and 5D) chromosomes. Therefore, it is unlikely that structural rearrangements could have a significant impact on the genomic patterns of introgression in the wheat-*Ae. tauschii* crosses.

**Fig. 3.**
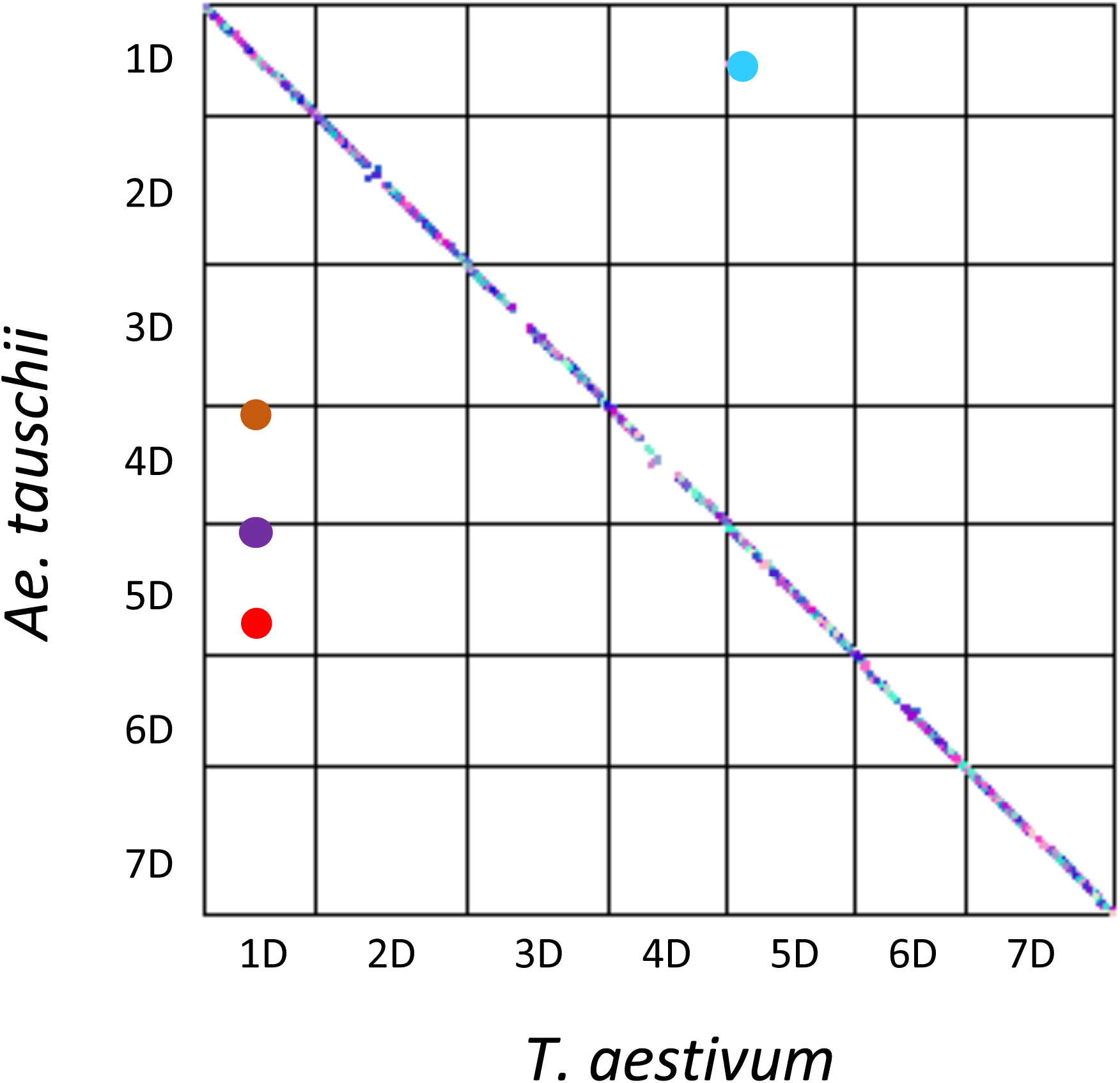
Gene collinearity and structural variation between the *T. aestivum* and *Ae. tauschii* chromosomes. Each dot on the dot-plot corresponds to a single gene and the big colored dots off diagonal represent the non-syntenic genes between genomes of the two species.

Interspecific introgression may also be influenced by the distribution of recombination rate along the chromosomes (Martin *et al.* 2019). Since strong depression of recombination in the pericentromeric regions creates steep recombination rate gradient along the centromere-telomere axis of large wheat chromosomes (Saintenac, Faure, *et al.* 2011; Jordan *et al.* 2018; Gardiner *et al.* 2019), we should expect distinct patterns of introgression in the pericentromeric and distal regions. To investigate the relationship between recombination and introgression, first, we calculated the number of recombination breakpoints (RBP) in 2 Mbp windows across the wheat genome across all families. Using the 95^th^ percentile of window-based RBP estimates as a threshold, we identified 200 genomic regions with a high recombination rate (Table 2, Table S4). To test whether the distribution of these regions with elevated recombination is random or associated with specific genomic regions, we compared their locations with the positions of high-recombining regions previously detected in the nested association mapping (NAM) population (Jordan *et al.* 2018) (Fig. 4a). We detected 43 overlapping regions with elevated recombination between these two populations (Table S4), mostly located in the distal chromosomal regions. This level of overlap was 6 times higher than the average overlap (7.2 RBPs) observed between the randomized datasets generated by permuting 2 Mbp windows in both introgression and NAM populations (Fig. 4b and 4c), suggestive of the presence of region-specific genomic features promoting recombination.

**Table 2.** Summary of 2 Mb windows recombination hotspots at 95^th^ percentile of total recombination breakpoints from 16 families of the introgression population.

| Chromosome | No. of windows | Min recombination | Max recombination |
| --- | --- | --- | --- |
| chr1D | 25 | 755 | 1968 |
| chr2D | 33 | 705 | 1441 |
| chr3D | 31 | 507 | 1136 |
| chr4D | 26 | 736 | 1499 |
| chr5D | 29 | 1052 | 1552 |
| chr6D | 24 | 899 | 1940 |
| chr7D | 32 | 869 | 1783 |

**Fig. 4.**
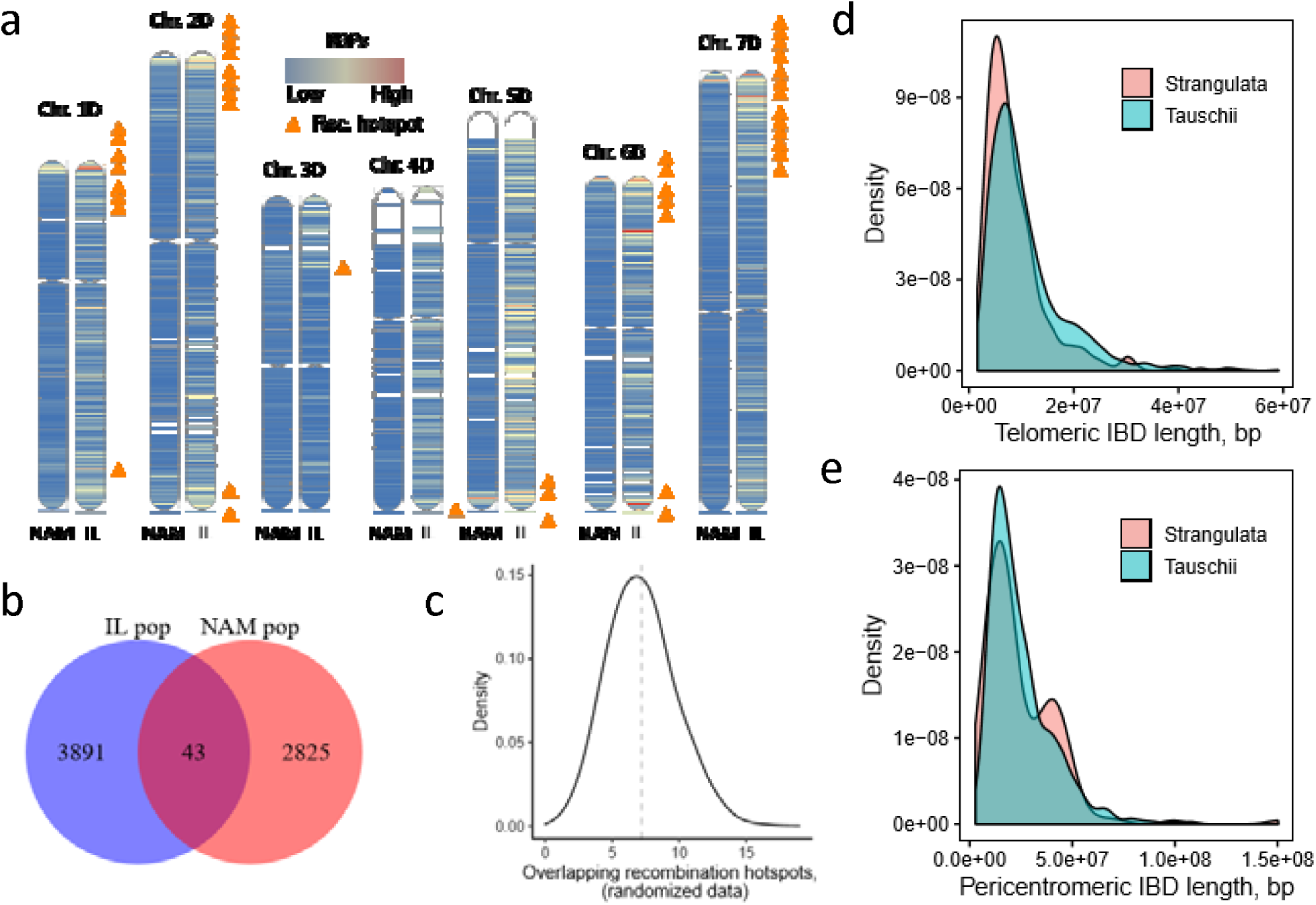
Effect of chromosome physical location on recombination rate and introgression. (a) A comparison of normalized recombination rate in 2 Mbp windows between nested association mapping (NAM) (Jordan et al. 2018) and *Ae. tauschii* introgression (IL) populations. The orange triangles show the overlapping windows with high recombination rate in the two populations at 95^th^ percentile. (b) A venn diagram showing the total number of 2 Mbp windows with at least one recombination event across the NAM and IL population while the intersection represents the overlaps at 95^th^ percentile. (c) Distribution of 2 Mbp window recombination hotspots permutation overlaps between the two populations. (d) Distribution of the introgressed segment’s length from *Ae. tauschii* ssp. *strangulata* and *Ae. tauschii* ssp. *tauschii* in the distal regions of the chromosomes. (e) Distribution of the introgressed segment’s length from *Ae. tauschii* ssp. *strangulata* and *Ae. tauschii* ssp. *tauschii* in the 2/3 pericentromeric regions of the chromosomes.

Second, we assessed the frequency of introgression along the wheat chromosomes. The IBD analysis was used to infer the boundaries of segments introgressed from *Ae. tauschii* into each line. A density plot of IBD segments along the chromosomes of the introgression population showed a U-shaped distribution (Fig. 5). The frequency of IBD segments positively correlated with the distribution of recombination rate (Jordan *et al.* 2018) and increased from the centromeres towards the telomeric regions of the chromosomes except on chromosome 1DL and 5DL in some families (Table S5). There was no chromosome preference during introgression. Variation in the number of introgressed segments per line was observed across chromosomes with the proportion of introgressed *Ae. tauschii* genome per line ranging from 0.05% to 25.5% (Table S6), deviating from the expected 25% of *Ae. tauschii* genome in the BC_1_F_3:5_ lines. Therefore, the efficiency of introgression in the population ranged from 0.2% to 100%. Some lines had single introgression while others had multiple introgressions per chromosome (Table S7). The IBD segments shared between the introgression lines and wheat parents were on average 5.9-fold longer than those shared with the *Ae. tauschii* parents (Table 3), and significantly different at the 95% confidence level based on the *t*-test statistics (P < 2.2e-16). The average length of IBD segments, expressed as a percent of the D genome size, shared between introgression lines and *Ae. tauschii* varied from 2.08% to 3.42% with a minimum of 0.25% and a maximum of 57.19%. Similarly, the average percent length of IBD segments shared between the chromosomes of introgression lines and hexaploid wheat parents ranged between 8.59% and 25.74% with a minimum of 0.36% and a maximum of 97.06%.

**Table 3.**
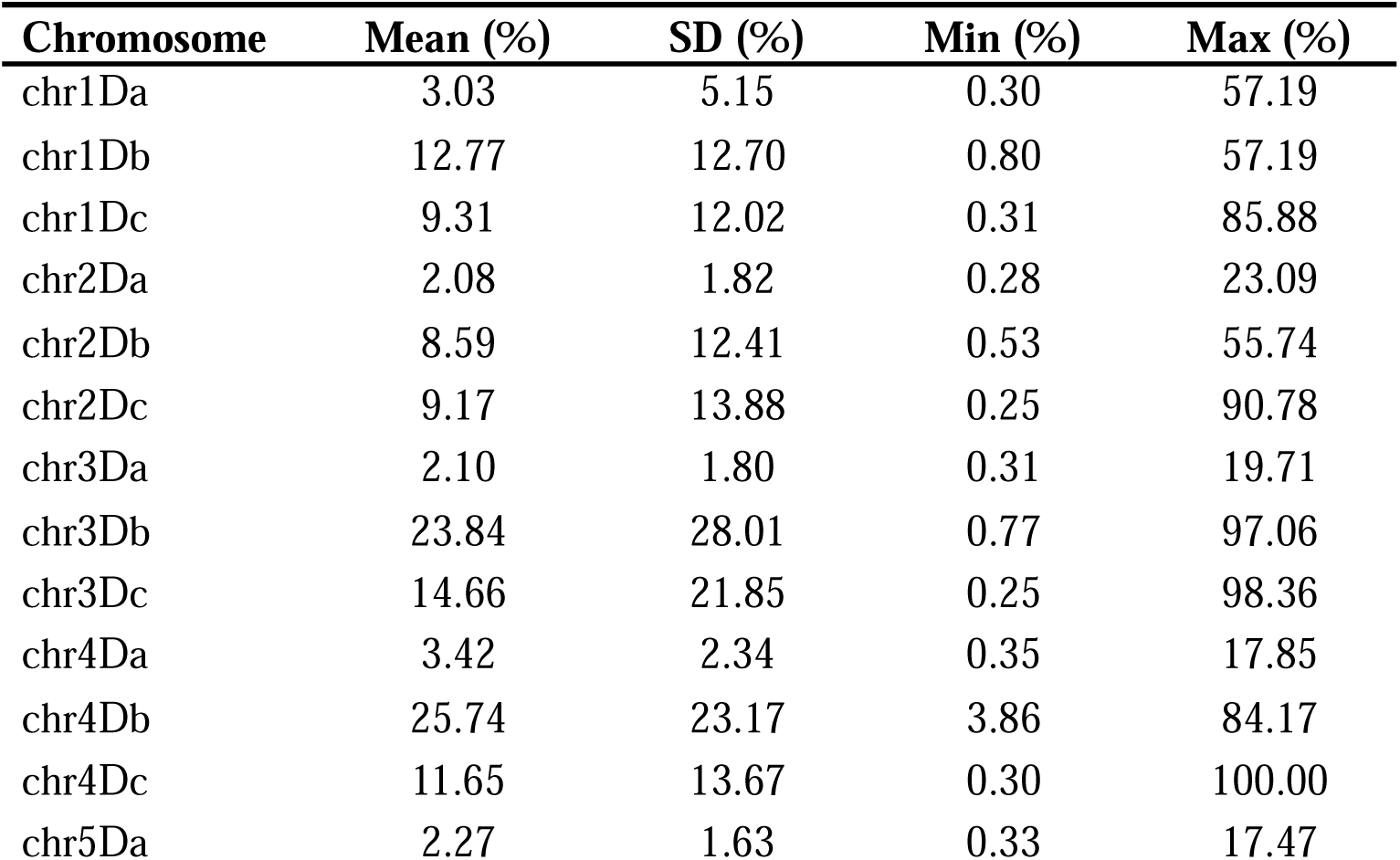
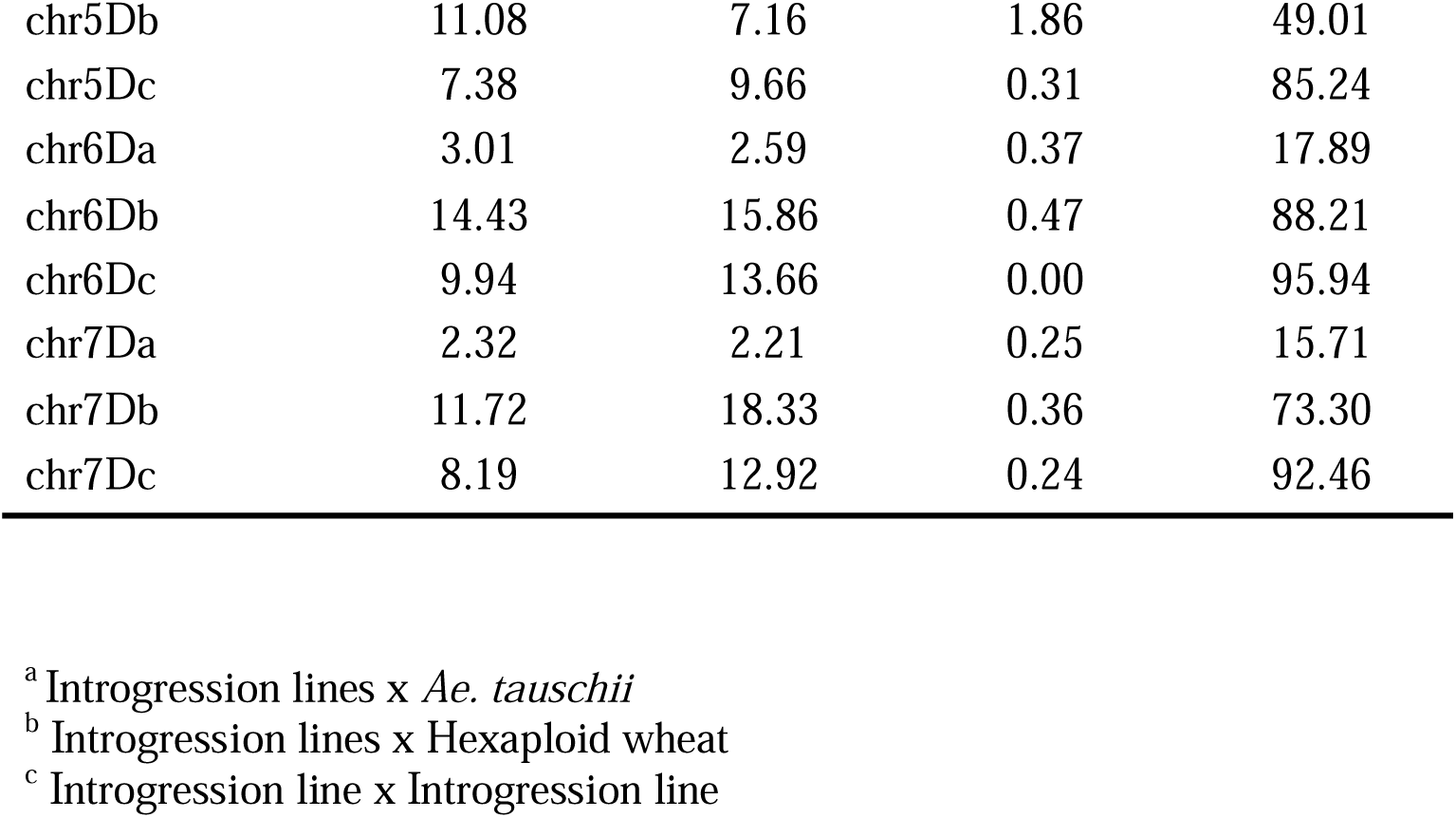
Percentage length of identity by descent segments shared between introgression lines, hexaploid wheat and *Ae. tauschii* accessions.

**Fig. 5.**
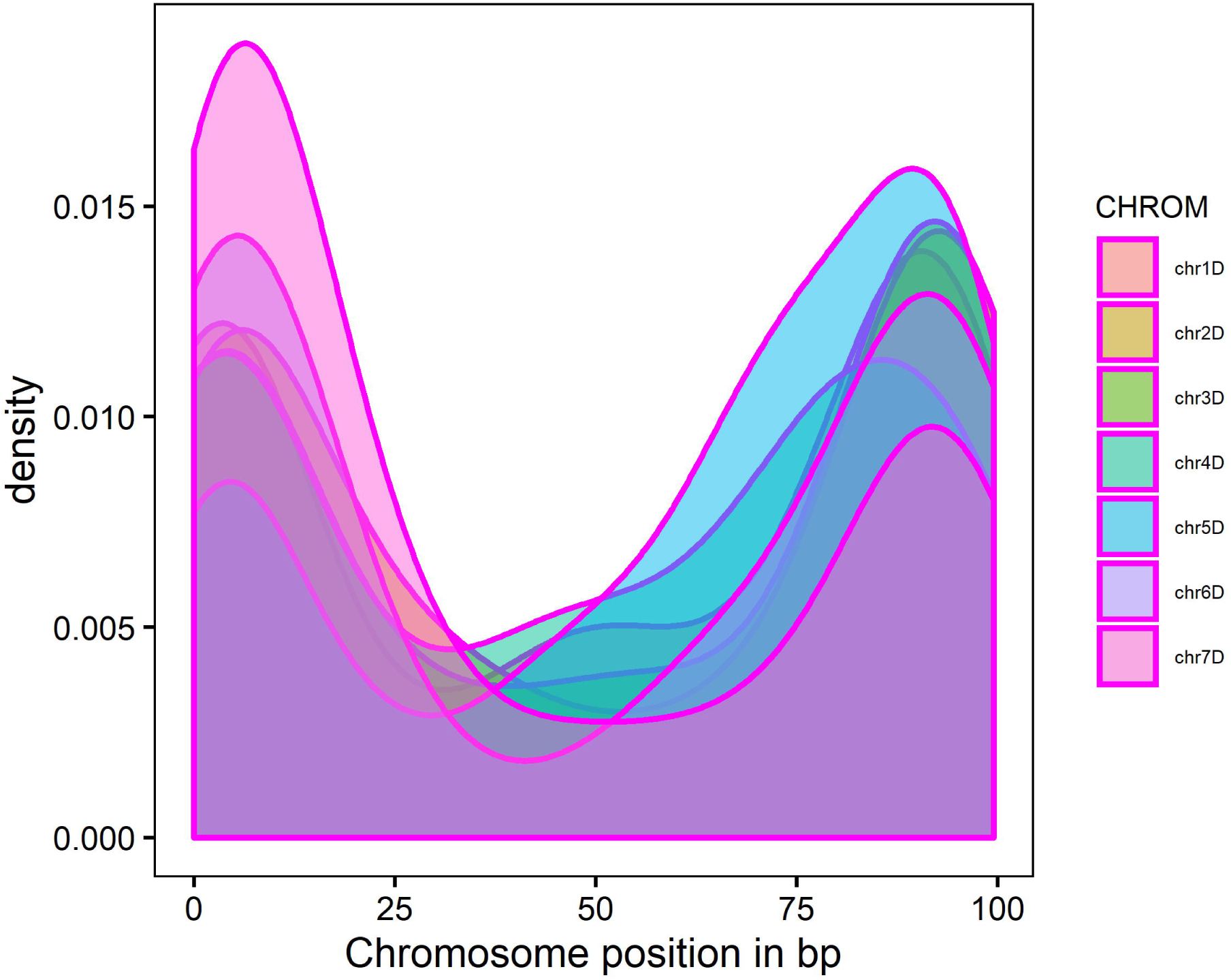
Density plots of identity by descent segments positions along the seven D genome chromosomes of the introgression lines derived from hexploid wheat and *Ae. tauschii* crosses. Chromosomes were scaled by dividing the IBD values by the individual chromosome length and then multiplied by 100.

### Relationship between sequence divergence, recombination and introgression length and frequency

The low rate of introgression in the pericentromeric regions may be the consequence of negative multilocus epistasis (Jiang et al. 2000; Martin et al. 2019) associated with introgression of large chromosomal segments. We compared the distribution of introgression segment’s length in the distal and pericentromeric regions of chromosomes between families derived from crosses with Ae. tauschii ssp. strangulata (direct donor of the wheat D genome), which shows a low level of divergence from the wheat D genome, and with Ae. tauschii ssp. tauschii, which shows higher levels of divergence from both the wheat D genome and Ae. tauschii ssp. strangulata (Figs. 4d and 4e) (Wang et al. 2013). No substantial differences in the length of introgressed segments were observed between the two subspecies in the distal regions. The mean length of introgressed segments was 8.8 Mbp (2.0 – 49.1 Mbp) for *Ae. tauschii ssp. strangulata* and 10.9 Mbp (1.6 – 59.1 Mbp) for *Ae. tauschii ssp. tauschii.* However, in the pericentromeric regions, we found an increase in the length of introgressed segments from *Ae. tauschii* ssp. *strangulata* (2.8 – 150 Mbp) compared to that from *Ae. tauschii* ssp. *tauschii* (3.9 – 107 Mbp) (Fig. 4e). Even though the mean length of introgression was not significantly different (24.5 Mbp and 24.7 Mbp, respectively) between the subspecies, *Ae. tauschii* ssp. *strangulata* showed bi-modal introgression size distribution, with a second density peak corresponding to introgressed segments with the mean of ∼40 Mbp (Fig. 4e).

We also compared the parental sequence divergence (SD) in the 2/3 pericentromeric and telomeric regions with IF between three *Ae. tauschii* ssp. *strangulata* derived families (FAM92, FAM93 and FAM96) and three *Ae. tauschii* ssp. *tauschii* derived families (FAM97, FAM98 and FAAM99) (Fig. S4). The mean SD was significantly different between the two subspecies with *ssp. strangulata* exhibiting a 4-fold lower SD (1.47 SNPs / 5 Mbp window) than *ssp. tauschii* (5.78 SNPs / 5 Mbp window) in the pericentromeric region with Mann-Whitney U test *p*-value < 2.2e-16. The low SD in the pericentromeric region of ssp. *strangulata* and hexaploid wheat parents was associated with a 2-fold increase in IF in the progenies. Similarly, SD and IF were significantly different in the telomeric regions of the two subspecies with Mann-Whitney U test *p*-value = 1.105e-15. The mean telomeric SD between hexaploid wheat and *Ae. tauschii* ssp. *strangulata* parents (10.6 SNPs / 5 Mbp window) was ∼2-folds lower than that between hexaploid wheat and *Ae. tauschii* ssp. *tauschii* parents (19.45 SNPs / 5 Mbp window). This resulted into a 1.3-fold increase in introgression frequency from ssp. *strangulata* (4.07) compared to that from ssp. *tauschii* (3.06). Taken together, these results suggest that the lower levels of divergence between parental genomes is linked with improved introgression efficiency, likely due to reduced negative multilocus epistasis.

The sequence divergence between the parental genomes might influence the frequency of homologous recombination (Opperman *et al.* 2004). In the pericentromeric regions, the mean number of RBPs estimated from ssp. *strangulata* (0.89) was 4-times lower than that estimated from ssp. *tauschii* derived families (3.44) (Fig. S4). Similarly, in the distal chromosomal regions, the estimated mean number of RBPs was higher in ssp. *tauschii* (5.94) than in ssp. *strangulata* (2.62) derived families. The high number of RBPs observed in ssp. *tauschii* compared to ssp. *strangulata* derived families may be explained by biases resulting from selection in the progenies of each family, and the low level of SNP diversity between the wheat D genome and *Ae. tauschii* ssp. *strangulata*, which is considered to be the donor of the wheat D genome (Dvorak *et al.* 1998), resulting in an underestimation of the total number of crossovers in the ssp. *strangulata* derived families. It is also possible that the increase in the levels of interhomolog polymorphism may stimulate recombination. In *Arabidopsis*, an increase in crossovers was observed when heterozygous regions were juxtaposed with homozygous regions (Ziolkowski *et al.* 2015), suggesting that the genomic distribution of interhomolog divergence have substantial effect on distribution of recombination rate.

### Patterns of introgression near the *Tg* locus

The free-threshing trait in wheat is controlled by the recessive variant of the *Tg* gene and the dominant variant of the *Q* gene (Simons *et al.* 2006; Sood *et al.* 2009; Dvorak *et al.* 2012; Faris *et al.* 2014). To understand the effect of selection against the hulled trait controlled by the dominant variant of *Tg* from *Ae. tauschii*, we investigated the patterns of IBD around the *Tg* locus shared by the introgression population with the *Ae. tauschii* parental genomes. We expected that the selection for mechanical seed threshing imposed during the introgression population development should result in a deficiency of the dominant *Tg* allele from *Ae. tauschii*, and increased frequency of the recessive *tg* allele contributed by the hexaploid wheat parents. To test this hypothesis, we identified the boundaries of the *Tg-D1* locus on chromosome 2D using two markers based on the expressed sequence tags (ESTs), CA658378 and BE518031 (Dvorak *et al.* 2012). These markers mapped to the reference wheat genome at positions 22,447,585 bp (BE518031) and 27,924,545 bp (CA658378). Additionally, the sequences of microsatellite markers *Xgwm*455, *Xgwm*296, *Xgwm*261 and *Xwmc*503 linked to *Tg* were aligned to the Chinese Spring reference v.1.0. Marker *Xwmc*503 closest to *Tg* gene mapped at 19.6 Mbp on 2DS (Table S8). Based on Sood *et al.* (2009) genetic map, the *Tg* gene is located 2.2 cM away from marker *Xwmc*503, implying that the *Tg* gene is located approximately at position 21.8 Mbp, which is consistent with the genomic interval flanked by markers based on CA658378 and BE518031 sequences. A count of IBD segments within 1-kbp sliding windows showed a steep decline in IBD between introgression lines and *Ae. tauschii* parents within the *Tg* gene region (Fig. 6a). This trend was accompanied by an increase in the proportion of IBD segments shared between the introgression lines and the hexaploid wheat parents, indicating a selection pressure for free-threshing trait during population development. The lowest decline in IBD segments count was observed around 22.8 Mbp. There are 40 high confidence genes within the 21.8 Mbp to 23.3 Mbp interval (Table S9), including two transcription factors from the bZIP and GRAS families.

**Fig. 6.**
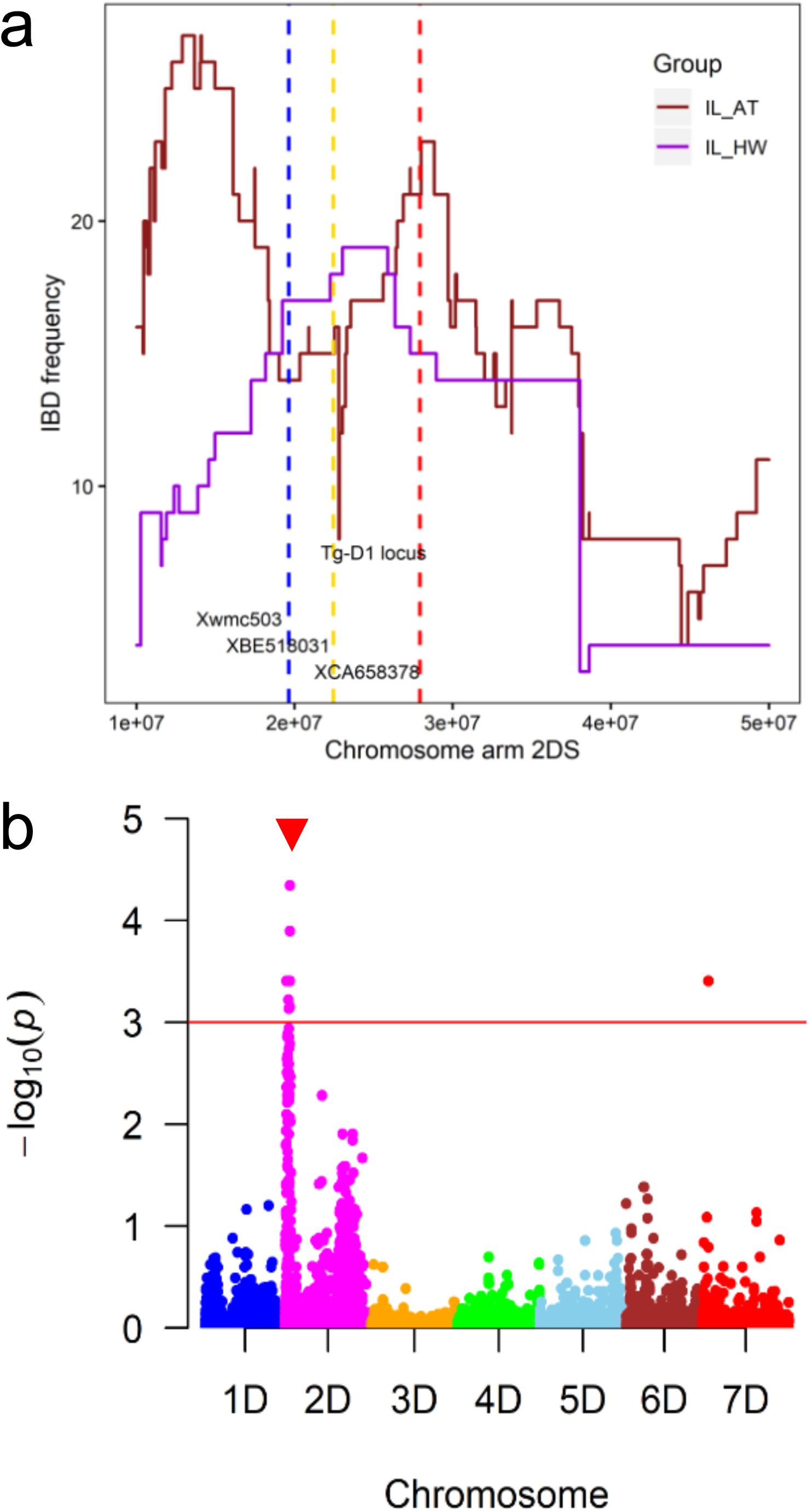
Location of *Tg* locus on chromosome arm 2DS as inferred by identity by descent (IBD) analysis and genome-wide association mapping. **a).** Shows the frequency of introgression from *Ae. tauschii* into hexploid wheat as inferred by IBD for chromosome arm 2DS region containing tenacious glume (*Tg*) gene. The IBD segments were counted per 1-kb sliding window. Where IL_AT and IL_HW are the trends of IBD frequency between introgression lines and *Ae. tauschii*, and hexaploid wheat, respectively. The blue line shows the position of marker *Xmwc503* (Sood et al. 2009), gold and red lines indicate the positions of EST markers *XBE518031* and *XCA658378*, respectively, flanking the *Tg-D1* locus. **b).** Manhattan plot showing the position of significant SNPs on 2DS (red arrow) and the red horizontal line shows the SNPs that are significant at an FDR q-value of 0.001.

To verify the impact of introgression on free-threshing, we phenotyped a subset of introgression lines for tenacious glume trait and compared the results with the IBD map (Fig. 6a). Most lines that had introgression spanning the *Tg* gene region were positive for tenacious glume trait (Table S10). Some lines that were shown to carry introgression at the *Tg* locus were not available for phenotyping because they were removed during harvesting by combine. As expected, lines showing no or small overlap of the introgressed segments with the *Tg* region were negative for tenacious glume trait.

Genome-wide association mapping was used to identify markers associated with the threshability trait. Using a mixed linear model while controlling for the population structure, we observed that the majority of the significant SNPs associated with tenacious glume trait in the introgression population were located on chromosome arm 2DS (Fig. 6b), which was consistent with the results of the IBD analysis. At a threshold FDR q-value of 0.05, 84 SNPs distributed along the *Tg* locus on 2DS showed a significant association with the trait, consistent with the previously reported maps (Table S11).

## Discussion

Loss of genetic diversity associated with domestication and improvement (Haudry *et al.* 2007; Akhunov *et al.* 2010; Ozkan *et al.* 2011; Xu *et al.* 2012; Hufford *et al.* 2012) can potentially reduce the adaptive potential of cultivated crops. Wild relatives found in regions with diverse agroecological environments are a valuable source of adaptive diversity that can improve resilience to biotic and abiotic stressors, or positively impact yield and quality traits (Uauy *et al.* 2006; Reynolds *et al.* 2007; Sohail *et al.* 2011; Cooper *et al.* 2012; Saintenac, Jiang, *et al.* 2013; Periyannan *et al.* 2013; Wulff and Moscou 2014; Chen *et al.* 2015). However, effective deployment of wild relative diversity in breeding requires a better understanding of the factors that represent a barrier for introgression into adapted germplasm. In this study, we used the progenies of crosses between bread wheat and lines carrying the genome of the diploid wild relative *Ae. tauschii* to investigate factors that have potential to impact the genomic patterns of introgression in the D genome of wheat.

As expected, we observed an increase in the number of polymorphic sites after introgression, with the increase being higher in the families derived from crosses involving *Ae. tauschii* ssp. *tauschii* rather than in the families derived from crosses with *Ae. tauschii* ssp. *strangulata*. The latter is consistent with the origin of the wheat D genome from *Ae. tauschii* ssp. *strangulata* lineage (Dvorak *et al.* 2012; Wang *et al.* 2013). In the families derived from both lineages of *Ae. tauschii*, we observed an uneven distribution of introgressed diversity along the chromosomes. The frequency of IBD regions shared with *Ae. tauschii* along all chromosomes showed a U-shaped distribution with lower incidence of introgression in the pericentromeric regions, indicating that the pericentromeric regions of chromosomes represent a more significant barrier to introgression than the distal chromosomal regions.

The introgression frequency correlated negatively with the length of IBD regions, sequence divergence between the parental genomes and positively with recombination rate, suggesting that longer introgressed segments in the low-recombining pericentromeric regions had a lower chance of being inherited in the progeny of crosses between *Ae. tauschii*-derived octoploids and wheat. These chromosomal patterns of introgression efficiency and length suggest that introgression was strongly affected by the distribution of recombination rate along chromosomes. This conclusion is consistent with recent research on butterflies that showed that the rate of inter-species introgression could be predicted by variation in recombination rate (Martin *et al.* 2019). It is likely that selection applied at BC_1_F_3:4_ generation to maintain uniform phenology, threshability, flowering time, and developmental characteristics inadvertently eliminated many lines carrying large introgressed regions in the pericentromeric regions. Our results also suggest that introgression from less divergent wild relative have lower probability of being eliminated from population that introgression from more divergent wild relative. Negative selection against introgression was previously observed in the interspecific crosses of polyploid cotton (Jiang *et al.* 2000), and attributed to multilocus negative epistasis between introgressed alleles and the genetic backgrounds of recipient lines. According to theory, introgressions that carry alleles having a negative impact on the selected traits will be removed from the population, with the size of the affected region defined by the recombination rate (Hill and Robertson 1966). It appears that a negative interaction between alleles located within the large introgressions in the low-recombining pericentromeric region and alleles of the adapted recurrent parent affected phenotypes targeted by selection, resulting in a loss of certain genotypes during population development. Thus, the limited number of recombination events at the BC_1_F_2_ generation, especially in the large pericentromeric regions of wheat chromosomes, resulted in linkage drag that affected a substantial proportion of the genome.

On the contrary, the terminal regions of the wheat chromosomes showed a high rate of introgression consistent with the theoretical predictions of the effect of selection on linked variation (Hill and Robertson 1966). In these regions, negatively selected alleles could be separated from the introgressed segments, as was clearly demonstrated for the *Tg* locus controlling free-threshing trait in wheat (Jantasuriyarat *et al.* 2004; Sood *et al.* 2009; Dvorak *et al.* 2012; Faris *et al.* 2014). Since this gene is located in the high-recombining terminal region of chromosome, we did not observe a substantial effect of selection against the wild-type allele on the frequency of introgression from *Ae. tauschii* outside of the *Tg* locus. The high recombination rate allowed for separating the negatively selected dominant *Tg* allele from the introgression and mapping the *Tg* gene locus to a 1.5 Mbp genomic interval, which was further confirmed by genome-wide association mapping.

Consistent with earlier studies in the field of speciation genomics (Martin and Jiggins 2017; Martin *et al.* 2019), barriers to introgression in wheat likely have polygenic nature with multiple negatively selected alleles affecting agronomic performance traits distributed across entire genome. As a result, the unintended consequence of selection applied at the early generation of interspecific crosses is the low rate of introgression in the large low-recombining regions of wheat chromosomes. This outcome suggests that multiple genes with a strong combined effect on adaptive traits are present in these regions, and identification of beneficial alleles in these regions will be complicated by linkage drag.

It is common practice for germplasm development programs to subject lines to selection pressure from early stages in population development. This approach is consistent with the goal of quickly identifying high performing lines to support commercial breeding, and allows for rapid exploitation of beneficial alleles in the chromosomal regions with high recombination. While this is a worthy objective, our results are a clear justification for a two-tiered approach to germplasm development aimed to maximize the exploitation of genetic diversity present in wild relatives. The first step is to ensure that maximum diversity is maintained in the introgression population. This could be achieved by applying low selection pressure and genotyping early generation populations to select subsets of lines carrying introgressions covering the genome. The second step is to maintain introgressed segments in the heterozygous state through random mating using approaches based on genetic male sterility or chemical hybridizing agents (Qi and Gill 2001; Singh *et al.* 2015; Guttieri 2019), that are warranted in self-pollinated species. This step would enhance effective recombination and increase the probability of separating beneficial alleles from the influence of linked negatively selected alleles in the regions of low recombination.

Recently, genetic factors controlling crossover rate across the genome and in the pericentromeric regions of wheat chromosomes have been identified (Jordan *et al.* 2018; Gardiner *et al.* 2019). The discovery of these genetic factors could also facilitate strategies to further increase the efficiency of introgression, and selection for favorable introgressed alleles in the low recombining regions. Failure to engage such strategies will result in the substantial loss of introgressed diversity across large portions of the genome, reducing the potential long-term impact of germplasm development programs.

## Acknowledgements

This project was supported by the Agriculture and Food Research Initiative Competitive Grant 2016-67013-24473 from the USDA National Institute of Food and Agriculture. We would like to thank Bliss Betzen for preparing genomic libraries, Alina Akhunova and KSU Integrated Genomics Facility for sequencing genomic libraries, and Jon Raupp from Wheat Genetics Resources Center for providing seeds of 21 accessions of *Aegilops tauschii* used for developing introgression population.

## Supporting information

### Supplemental tables

Table S1. Summary of GBS data for introgression lines, *Ae. tuaschii* and hexaploid wheat parents.

Table S2. Summary of GBS data for 116 *Ae. tauschii* accessions used as a reference panel. Table S3. Origin of *Ae. tauschii* accessions used as reference panel, the source of 21 *Ae. tauschii* used as introgression parents and their grouping based on the first two principal components.

Table S4. Frequency of total recombination breakpoint from 16 introgression population families.

Table S5. Pearson’s correlation coefficients between recombination rate and introgression frequency within 2 Mbp windows, and between recombination rate and the physical genomic distance from the centromere toward the telomeres on both chromosome arms in 16 *Ae. tauschii* introgression families.

Table S6. Efficiency of *Ae. tauschii* introgression in wheat as inferred by identity by descent.

Table S7. Location of introgressed genomic segments with the highest LOD values from *Ae. tauschii* in hexaploid wheat background as predicted by Beagle v4.1.

Table S8. Location of microsatellite and expressed sequence tags (EST) markers linked to tenacious glume (*Tg*) gene on the Chinese Spring reference v1.0.

Table S9. High confidence genes within chromosome arm 2DS interval known to control tenacious glume trait.

Table S10. Tenacious glume scores for the introgression lines with and without introgression from *Ae. tauschii* parents on chromosome arm 2DS where the *Tg* gene is located.

Table S11. SNPs on chromosome arm 2DS significantly associated with tenacious glume trait.

### Supplemental figures

Fig. S1. Distribution of 116 *Ae. tauschii* accessions (red, AT) used as reference panel and the 21 *Ae. tauschii* accessions (magenta, ILP_AT) used to generate the introgression lines on the first two principal components. L1a and L1b accessions belong to *Ae. tauschii* ssp. *tauschii* while L2 accessions belong to *Ae. tauschii* ssp. *strangulata*.

Fig. S2. Distribution of 116 *Ae. tauschii* accessions (AT) used as reference panel, the 21 *Ae. tauschii* accessions (ILP_AT) used to generate the introgression lines, the 6 hexaploid wheat parents (ILP_HW) and the 351 introgression lines (IL) on the first two principal components. L1a and L1b accessions belong to *Ae. tauschii* ssp. *tauschii* while L2 accessions belong to *Ae. tauschii* ssp. *strangulata*.

Fig. S3. Variation in nucleotide diversity per chromosome based on π values for *Ae. Tauschii* (AT) accessions, hexaploid wheat (HW) and introgression lines (IL).

Fig. S4. Effect of parental sequence divergence (SD) in the 2/3 pericentromeric and telomeric chromosomal regions on recombination rate (RBP) and introgression frequency (IF) in *Ae. tauschii* ssp. *strangulata* and *Ae. tauschii* ssp. *tauschii* derived introgression families.

## Notes

### Competing Interest Statement

The authors have declared no competing interest.

## References

Akhunov, E. D., A. R. Akhunova, O. D. Anderson, J. a Anderson, N. Blake et al., 2010 Nucleotide diversity maps reveal variation in diversity among wheat genomes and chromosomes. BMC Genomics 11: 702.

Avni, R., M. Nave, O. Barad, K. Baruch, S. O. Twardziok et al., 2017 Wild emmer genome architecture and diversity elucidate wheat evolution and domestication. Science 97: 93–97.

Balfourier, F., S. Bouchet, S. Robert, R. DeOliveira, H. Rimbert et al., 2019 Worldwide phylogeography and history of wheat genetic diversity. Sci. Adv. 5:.

Browning, S. R., and B. L. Browning, 2012 Identity by descent between distant relatives: detection and applications. Annu. Rev. Genet. 46: 617–33.

Browning, B. L., and S. R. Browning, 2013 Improving the accuracy and efficiency of identity-by-descent detection in population data. Genetics 194: 459–71.

Chen, S., M. N. Rouse, W. Zhang, Y. Jin, E. Akhunov et al., 2015 Fine mapping and characterization of Sr21, a temperature-sensitive diploid wheat resistance gene effective against the Puccinia graminis f. sp. tritici Ug99 race group. Theor. Appl. Genet. 128: 645–56.

Cooper, J. K., a. M. H. Ibrahim, J. Rudd, S. Malla, D. B. Hays et al., 2012 Increasing hard winter wheat yield potential via synthetic wheat: I. path-coefficient analysis of yield and its components. Crop Sci. 52: 2014–2022.

Dale, Z., H. Jie, H. Luyu, Z. Cancan, Z. Yun et al., 2017 An advanced backcross population through synthetic octaploid wheat as a “Bridge”: Development and QTL detection for seed dormancy. Front. Plant Sci. 8: 1–10.

Dreisigacker, S., M. Kishii, J. Lage, and M. Warburton, 2008 Use of synthetic hexaploid wheat to increase diversity for CIMMYT bread wheat improvement. Aust. J. Agric. Res. 59: 413–420.

Dvorak, J., K. R. Deal, M.-C. Luo, F. M. You, K. von Borstel et al., 2012 The origin of spelt and free-threshing hexaploid wheat. J. Hered. 103: 426–41.

Dvorak, J., M. C. Luo, Z. L. Yang, and H. B. Zhang, 1998 The structure of the Aegilops tauschii genepool and the evolution of hexaploid wheat. Theor. Appl. Genet. 97: 657–670.

Elshire, R. J., J. C. Glaubitz, Q. Sun, J. a Poland, K. Kawamoto et al., 2011 A robust, simple genotyping-by-sequencing (GBS) approach for high diversity species. PLoS One 6: e19379.

Faris, J. D., Q. Zhang, S. Chao, Z. Zhang, and S. S. Xu, 2014 Analysis of agronomic and domestication traits in a durum × cultivated emmer wheat population using a high-density single nucleotide polymorphism-based linkage map. Theor. Appl. Genet. 127: 2333–48.

Gardiner, L., L. U. Wingen, P. Bailey, R. Joynson, T. Brabbs et al., 2019 Analysis of the recombination landscape of hexaploid bread wheat reveals genes controlling recombination and gene conversion frequency. Genome Biol. 20: 69.

Gill, B. S., and W. J. Raupp, 1987 Direct Genetic Transfers from Aegilops squarrosa L. to Hexaploid Wheat. Crop Sci. 27: 445–450.

Glaubitz, J. C., T. M. Casstevens, F. Lu, J. Harriman, R. J. Elshire et al., 2014 TASSEL-GBS: A High Capacity Genotyping by Sequencing Analysis Pipeline. PLoS One 9: e90346.

Guttieri, M. J., 2019 Ms3 dominant genetic male sterility for wheat improvement with molecular breeding. Crop Sci. doi: 10.2135/cropsci2019.06.0378.

Haudry, a, a Cenci, C. Ravel, T. Bataillon, D. Brunel et al., 2007 Grinding up wheat: a massive loss of nucleotide diversity since domestication. Mol. Biol. Evol. 24: 1506–17.

He, F., R. Pasam, F. Shi, S. Kant, G. Keeble-Gagnere et al., 2019 Exome sequencing highlights the role of wild relative introgression in shaping the adaptive landscape of the wheat genome. Nat. Genet. 51: 896–904.

Hill, W., and Robertson, 1966 The effect of linkage on limits to artificial selection. Genet. Res. 8: 269–294.

Huang, L., S. A. Brooks, W. Li, J. P. Fellers, H. N. Trick et al., 2003 Map-based cloning of leaf rust resistance gene Lr21 from the large and polyploid genome of bread wheat. Genetics 164: 655–64.

Hufford, M. B., X. Xu, J. van Heerwaarden, T. Pyhäjärvi, J.-M. Chia et al., 2012 Comparative population genomics of maize domestication and improvement. Nat. Genet. 44: 808–11.

Jantasuriyarat, C., M. I. Vales, C. J. W. Watson, and O. Riera-Lizarazu, 2004 Identification and mapping of genetic loci affecting the free-threshing habit and spike compactness in wheat (Triticum aestivum L.). Theor. Appl. Genet. 108: 261–273.

Jiang, C. X., P. W. Chee, X. Draye, P. L. Morrell, C. W. Smith et al., 2000 Multilocus interactions restrict gene introgression in interspecific populations of polyploid Gossypium (cotton). Evolution (N. Y). 54: 798–814.

Jordan, K. W., S. Wang, F. He, S. Chao, Y. Lun et al., 2018 The genetic architecture of genome-wide recombination rate variation in allopolyploid wheat revealed by nested association mapping. Plant J. 95: 1039–1054.

Jordan, K., S. Wang, Y. Lun, L. Gardiner, R. MacLachlan et al., 2015 A haplotype map of allohexaploid wheat reveals distinct patterns of selection on homoeologous genomes. Genome Biol. 16: 48.

Kihara, H., 1944 Discovery of the DD-analyser, one of the ancestors of Triticum vulgare. Agricuture Hortic. 19: 889–890.

Ling, H.-Q., S. Zhao, D. Liu, J. Wang, H. Sun et al., 2013 Draft genome of the wheat A-genome progenitor Triticum urartu. Nature 496: 87–90.

Luo, M.-C., Y. Q. Gu, D. Puiu, H. Wang, S. O. Twardziok et al., 2017 Genome sequence of the progenitor of the wheat D genome Aegilops tauschii. Nature 551: 498–502.

Luo, M.-C., Z.-L. Yang, F. M. You, T. Kawahara, J. G. Waines et al., 2007 The structure of wild and domesticated emmer wheat populations, gene flow between them, and the site of emmer domestication. Theor. Appl. Genet. 114: 947–59.

Martin, S. H., J. W. Davey, C. Salazar, and C. D. Jiggins, 2019 Recombination rate variation shapes barriers to introgression across butterfly genomes. PLoS Biol. 17: 1–28.

Martin, S. H., and C. D. Jiggins, 2017 Interpreting the genomic landscape of introgression. Curr. Opin. Genet. Dev. 47: 69–74.

Neph, S., M. S. Kuehn, A. P. Reynolds, E. Haugen, R. E. Thurman et al., 2012 BEDOPS: High-performance genomic feature operations. Bioinformatics 28: 1919–1920.

Nyine, M., B. Uwimana, N. Blavet, E. Hřibová, H. Vanrespaille et al., 2018 Genomic Prediction in a Multiploid Crop: Genotype by Environment Interaction and Allele Dosage Effects on Predictive Ability in Banana. Plant Genome 11: 10.3835/plantgenome2017.10.0090.

Nyine, M., S. Wang, K. Kiani, K. Jordan, S. Liu et al., 2019 Genotype Imputation in Winter Wheat Using First-Generation Haplotype Map SNPs Improves Genome-Wide Association Mapping and Genomic Prediction of Traits. G3: Genes|Genomes|Genetics 9: 125–133.

Ogbonnaya, F. C., O. Abdalla, A. Mujeeb-Kazi, A. G. Kazi, S. S. Xu et al., 2013 Synthetic Hexaploids: Harnessing Species of the Primary Gene Pool for Wheat Improvement. Plant Breed. Rev. 35–122.

Opperman, R., E. Emmanuel, and A. A. Levy, 2004 The effect of sequence divergence on recombination between direct repeats in arabidopsis. Genetics 168: 2207–2215.

Ozkan, H., G. Willcox, A. Graner, F. Salamini, and B. Kilian, 2011 Geographic distribution and domestication of wild emmer wheat (Triticum dicoccoides). Genet. Resour. Crop Evol. 58: 11–53.

Peleg, Z., T. Fahima, S. Abbo, T. Krugman, E. Nevo et al., 2005 Genetic diversity for drought resistance in wild emmer wheat and its ecogeographical associations. Plant, Cell Environ. 28: 176–191.

Periyannan, S., J. Moore, M. Ayliffe, U. Bansal, X. Wang et al., 2013 The Gene Sr33, an Ortholog of Barley Mla Genes, Encodes Resistance to Wheat Stem Rust Race Ug99. Science 341: 786–8.

Poland, J. A., P. J. Brown, M. E. Sorrells, and J.-L. Jannink, 2012 Development of high-density genetic maps for barley and wheat using a novel two-enzyme genotyping-by-sequencing approach. PLoS One 7: e32253.

Pont, C., T. Leroy, M. Seidel, A. Tondelli, W. Duchemin et al., 2019 Tracing the ancestry of modern bread wheats. Nat. Genet. 51: 905–911.

Qi, L., B. Friebe, P. Zhang, and B. S. Gill, 2007 Homoeologous recombination, chromosome engineering and crop improvement. Chromosom. Res. 15: 3–19.

Qi, L. L., and B. S. Gill, 2001 High-density physical maps reveal that the dominant male-sterile gene Ms3 is located in a genomic region of low recombination in wheat and is not amenable to map-based cloning. Theor. Appl. Genet. 103: 998–1006.

R Development Core Team, R., 2011 R: A Language and Environment for Statistical Computing (R. D. C. Team, Ed.). R Found. Stat. Comput. 1: 409.

Reynolds, M., F. Dreccer, and R. Trethowan, 2007 Drought-adaptive traits derived from wheat wild relatives and landraces. J. Exp. Bot. 58: 177–86.

Reynolds, M. P., A. J. D. Pask, W. J. E. Hoppitt, K. Sonder, S. Sukumaran et al., 2017 Strategic crossing of biomass and harvest index—source and sink—achieves genetic gains in wheat. Euphytica 213: 213–257.

Rieseberg, L. H., C. R. Linder, and G. J. Seiler, 1995 Chromosomal and genic barriers to introgression in helianthus. Genetics 141: 1163–1171.

Rieseberg, L. H., J. Whitton, and K. Gardner, 1999 Hybrid zones and the genetic architecture of a barrier to gene flow between two sunflower species. Genetics 152: 713–727.

Saintenac, C., S. Faure, A. Remay, F. Choulet, C. Ravel et al., 2011 Variation in crossover rates across a 3-Mb contig of bread wheat (Triticum aestivum) reveals the presence of a meiotic recombination hotspot. Chromosoma 120: 185–198.

Saintenac, C., D. Jiang, and E. D. Akhunov, 2011 Targeted analysis of nucleotide and copy number variation by exon capture in allotetraploid wheat genome. Genome Biol. 12: R88.

Saintenac, C., D. Jiang, S. Wang, and E. Akhunov, 2013 Sequence-based mapping of the polyploid wheat genome. G3 (Bethesda). 3: 1105–14.

Saintenac, C., W. Zhang, A. Salcedo, M. N. Rouse, H. N. Trick et al., 2013 Identification of wheat gene Sr35 that confers resistance to Ug99 stem rust race group. Science 341: 783–6.

Sears, E. R., 1977 AN INDUCED MUTANT WITH HOMOEOLOGOUS PAIRING IN COMMON WHEAT. Can. J. Genet. Cytol. 19: 585–593.

Simons, K. J., J. P. Fellers, H. N. Trick, Z. Zhang, Y.-S. Tai et al., 2006 Molecular characterization of the major wheat domestication gene Q. Genetics 172: 547–55.

Singh, S., R. Srivastava, and J. Kumar, 2015 Male sterility systems in wheat and opportunities for hybrid wheat development. Acta Physiol Plant. 37: 1–13.

Singh, N., S. Wu, V. Tiwari, S. Sehgal, J. Raupp et al., 2019 Genomic analysis confirms population structure and identifies inter-lineage hybrids in Aegilops tauschii. Front. Plant Sci. 10: 1–13.

Sohail, Q., T. Inoue, H. Tanaka, A. E. Eltayeb, Y. Matsuoka et al., 2011 Applicability of Aegilops tauschii drought tolerance traits to breeding of hexaploid wheat. Breed. Sci. 61: 347–57.

Sood, S., V. Kuraparthy, G. Bai, and B. S. Gill, 2009 The major threshability genes soft glume (sog) and tenacious glume (Tg), of diploid and polyploid wheat, trace their origin to independent mutations at non-orthologous loci. Theor. Appl. Genet. 119: 341–51.

Su, H., Y. Liu, C. Liu, Q. Shi, Y. Huang et al., 2019 Centromere Satellite Repeats Have Undergone Rapid Changes in Polyploid Wheat Subgenomes. Plant Cell 31: tpc.00133.2019.

The International Wheat Genome Sequencing Consortium (IWGSC), 2018 Shifting the limits in wheat research and breeding using a fully annotated reference genome. Science 361: eaar7191.

Uauy, C., A. Distelfeld, T. Fahima, A. Blechl, and J. Dubcovsky, 2006 A NAC Gene Regulating Senescence Improves Grain Protein, Zinc, and Iron Content in Wheat. Science 314: 1298–1301.

Wang, L., T. M. Beissinger, A. Lorant, C. Ross-Ibarra, J. Ross-Ibarra et al., 2017 The interplay of demography and selection during maize domestication and expansion. Genome Biol. 18: 1–13.

Wang, J., M.-C. Luo, Z. Chen, F. M. You, Y. Wei et al., 2013 Aegilops tauschii single nucleotide polymorphisms shed light on the origins of wheat D-genome genetic diversity and pinpoint the geographic origin of hexaploid wheat. New Phytol. 198: 925–937.

Wang, Y., H. Tang, J. D. Debarry, X. Tan, J. Li et al., 2012 MCScanX: A toolkit for detection and evolutionary analysis of gene synteny and collinearity. Nucleic Acids Res. 40: 1–14.

Wulff, B. B. H., and M. J. Moscou, 2014 Strategies for transferring resistance into wheat: from wide crosses to GM cassettes. Front. Plant Sci. 5: 692.

Xu, X., X. Liu, S. Ge, J. D. Jensen, F. Hu et al., 2012 Resequencing 50 accessions of cultivated and wild rice yields markers for identifying agronomically important genes. Nat. Biotechnol. 30: 105–11.

Ziolkowski, P. A., L. E. Berchowitz, C. Lambing, N. E. Yelina, X. Zhao et al., 2015 Juxtaposition of heterozygous and homozygous regions causes reciprocal crossover remodelling via interference during Arabidopsis meiosis. Elife 4: 1–29.

