## Supplementary figures and images for "Genomic patterns of introgression in interspecific populations created by crossing wheat with its wild relative"

### Figure S1

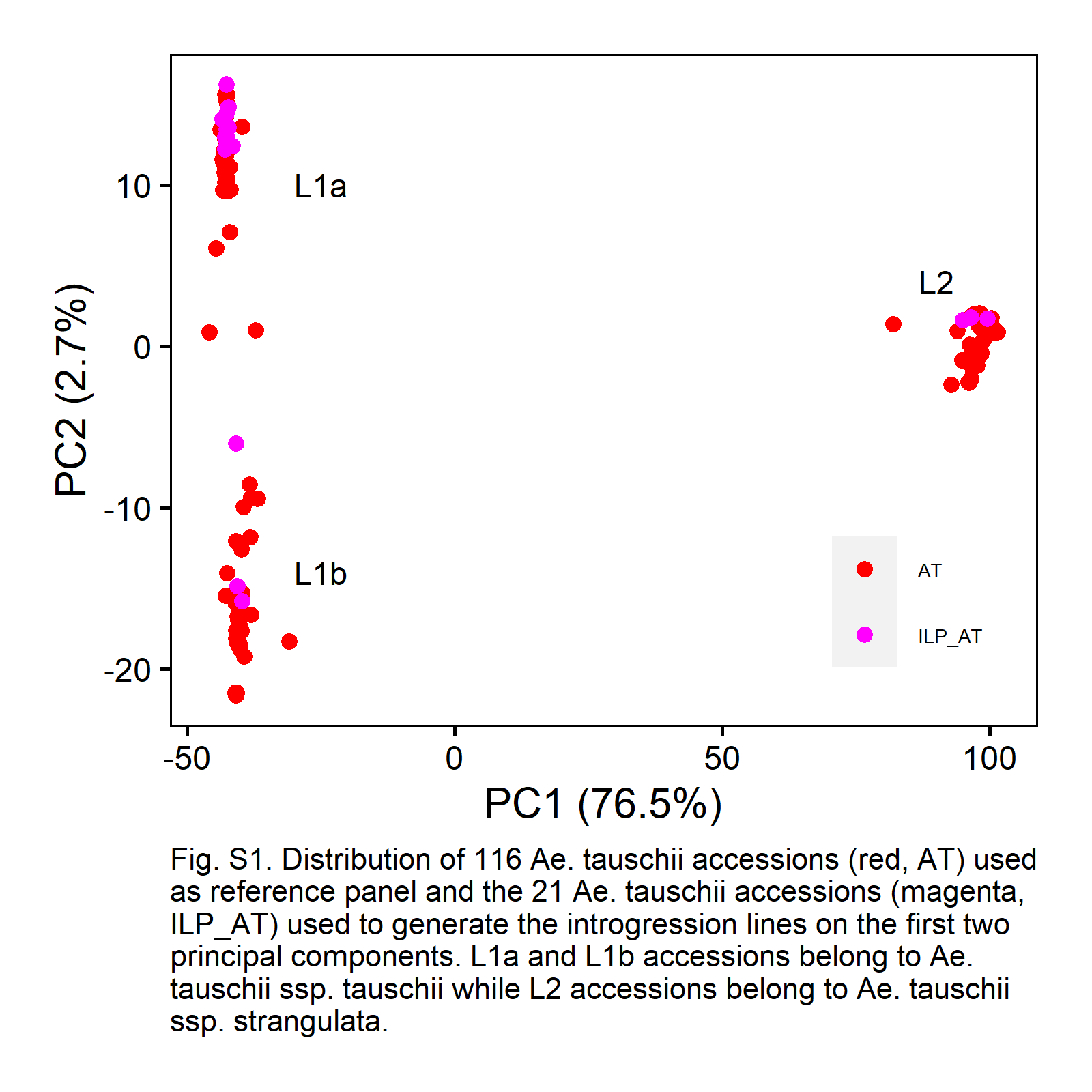

### Figure S2

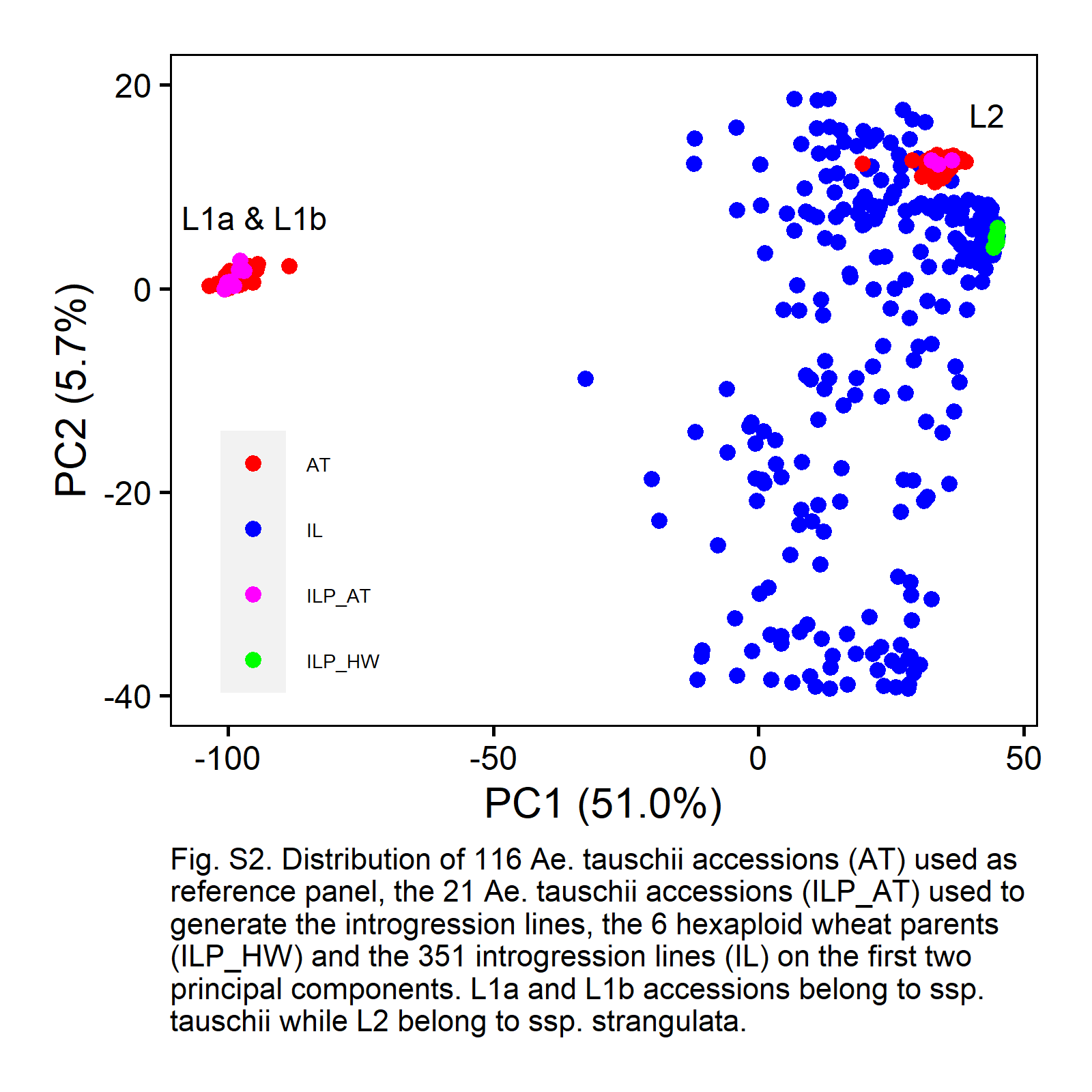
