## Supplementary material for "Genomic patterns of introgression in interspecific populations created by crossing wheat with its wild relative": Figure S3

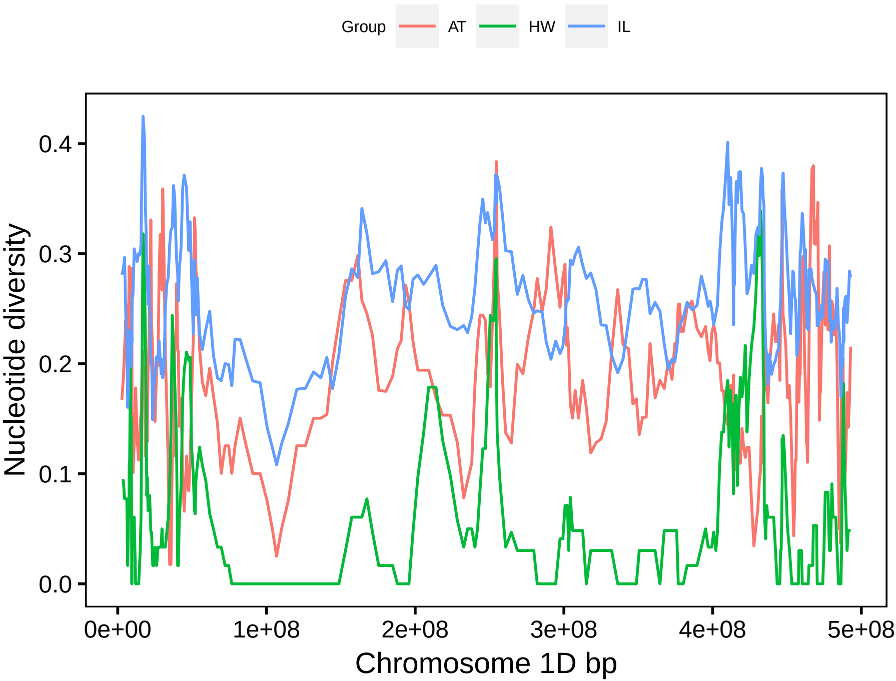

Group
AT
HW
IL
Nucleotide diversity
0.0
0.1
0.2
0.3
0.4
0e+00
1e+08
2e+08
3e+08
4e+08
5e+08
Chromosome 1D bp

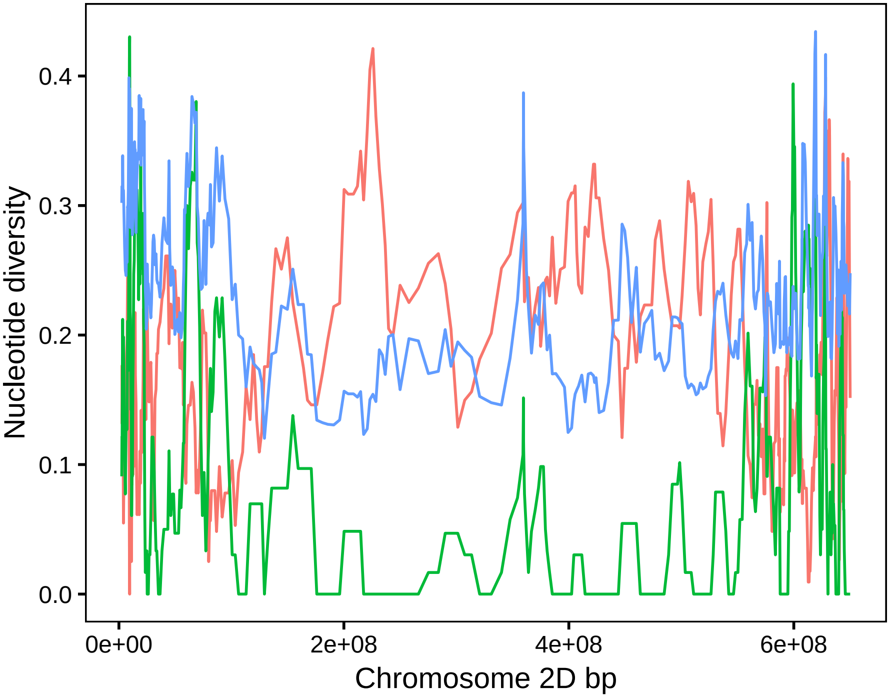

Nucleotide diversity
0.0
0.1
0.2
0.3
0.4
0e+00
2e+08
4e+08
6e+08
Chromosome 2D bp

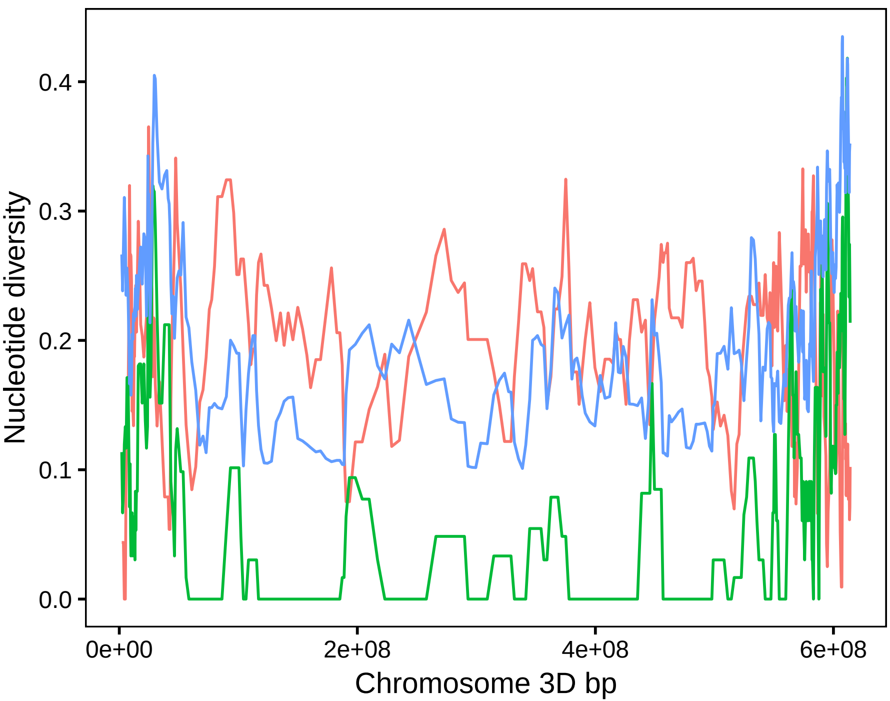

0.0
0.1
0.2
0.3
0.4
Nucleotide diversity
0e+00
2e+08
4e+08
6e+08
Chromosome 3D bp

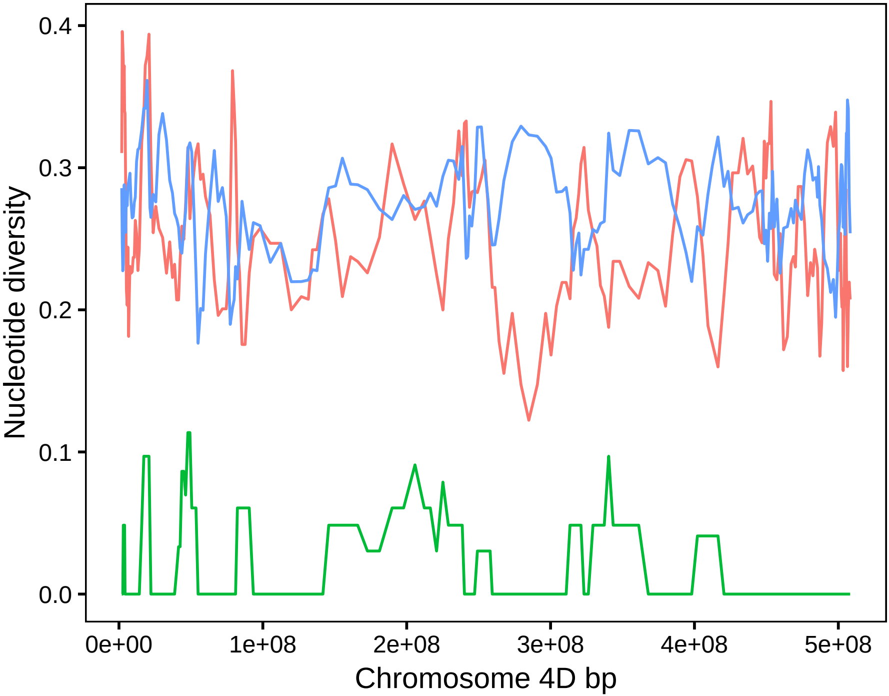
0.4
0.3
0.2
0.1
0.0
Nucleotide diversity
0e+00
1e+08
2e+08
3e+08
4e+08
5e+08
Chromosome 4D bp

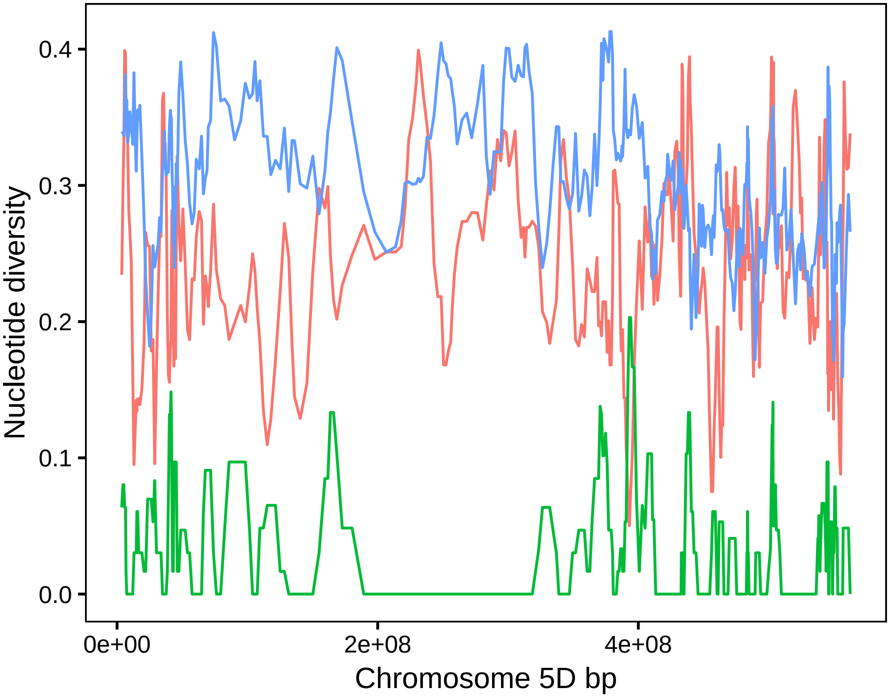

Nucleotide diversity
0.0
0.1
0.2
0.3
0.4
0e+00
2e+08
4e+08
Chromosome 5D bp

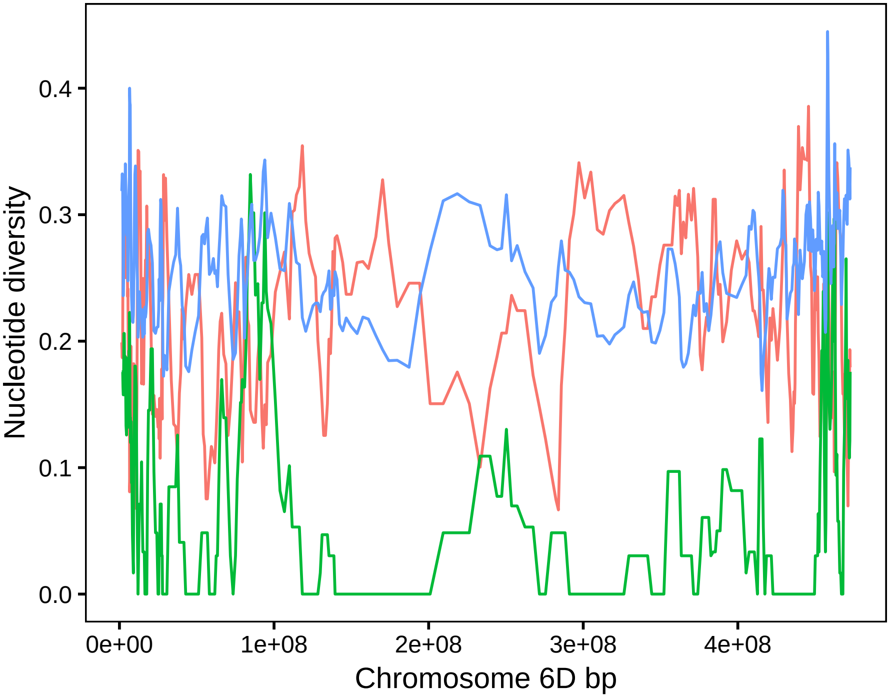

Nucleotide diversity
0.4
0.3
0.2
0.1
0.0
0e+00
1e+08
2e+08
3e+08
4e+08
Chromosome 6D bp

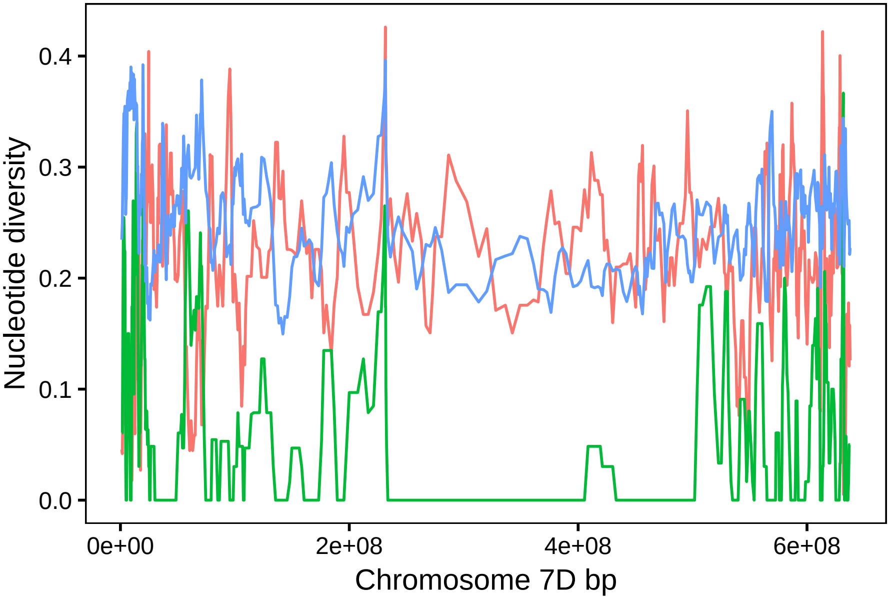


Fig. S3. Variation in nucleotide diversity per chromosome based on SNP pi values for Ae. tauschii (AT) accessions, hexaploid wheat (HW) and introgression lines (IL).
