## Supplementary material for "Genomic patterns of introgression in interspecific populations created by crossing wheat with its wild relative": Figure S4

### Slide 1
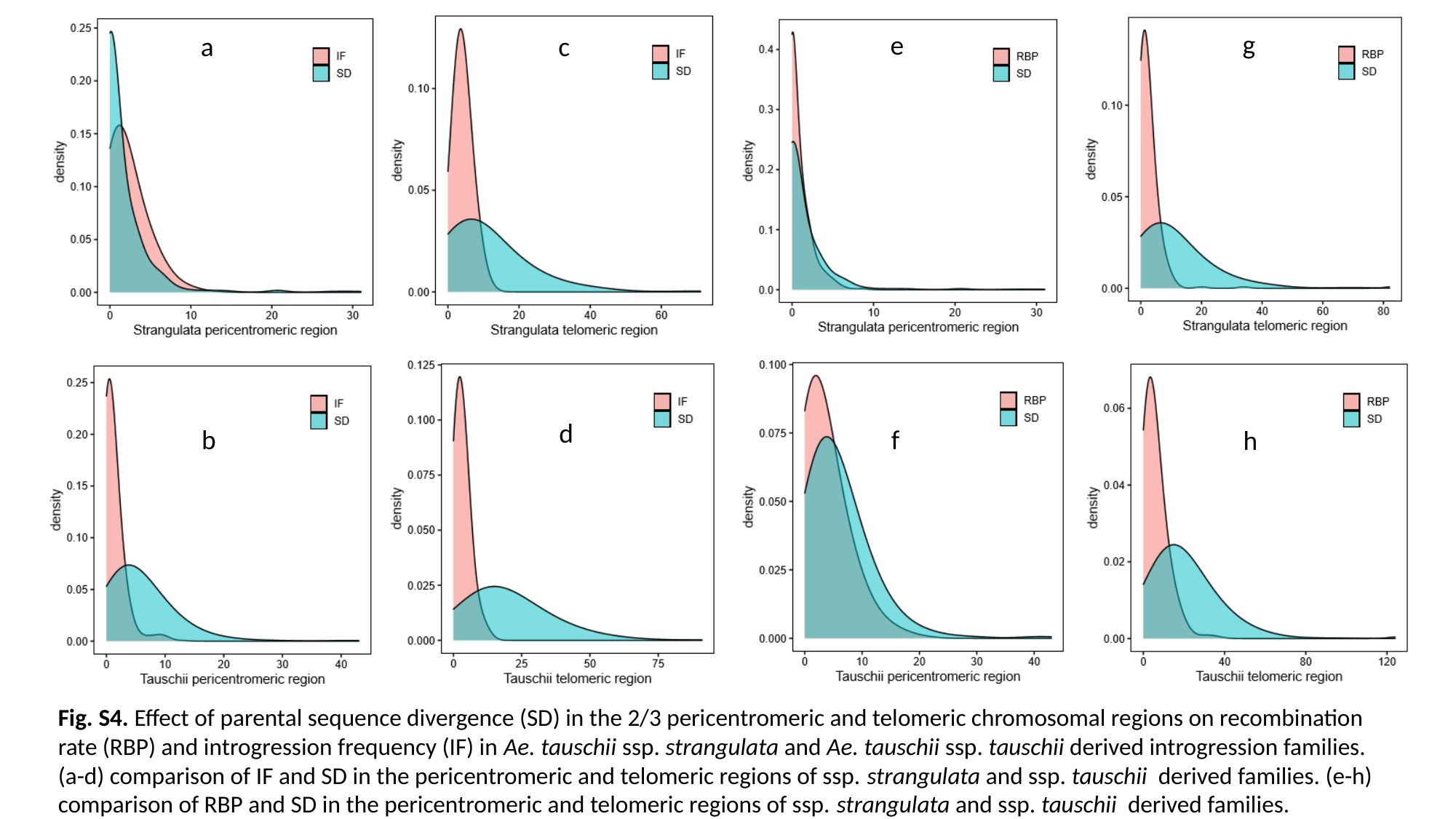

g
e
a
c
d
f
b
h
Fig. S4. Effect of parental sequence divergence (SD) in the 2/3 pericentromeric and telomeric chromosomal regions on recombination rate (RBP) and introgression frequency (IF) in Ae. tauschii ssp. strangulata and Ae. tauschii ssp. tauschii derived introgression families. (a-d) comparison of IF and SD in the pericentromeric and telomeric regions of ssp. strangulata and ssp. tauschii derived families. (e-h) comparison of RBP and SD in the pericentromeric and telomeric regions of ssp. strangulata and ssp. tauschii derived families.
